# Allele ages provide limited information about the strength of negative selection

**DOI:** 10.1101/2024.08.06.606888

**Authors:** Vivaswat Shastry, Jeremy J. Berg

## Abstract

For many problems in population genetics, it is useful to characterize the distribution of fitness effects (DFE) of *de novo* mutations among a certain class of sites. A DFE is typically estimated by fitting an observed site frequency spectrum (SFS) to an expected SFS given a hypothesized distribution of selection coefficients and demographic history. The development of tools to infer gene trees from haplotype alignments, along with ancient DNA resources, provides us with additional information about the frequency trajectories of segregating mutations. Here, we ask how useful this additional information is for learning about the DFE, using the joint distribution on allele frequency and age to summarize information about the trajectory. To this end, we introduce an accurate and efficient numerical method for computing the density on the age of a segregating variant found at a given sample frequency, given the strength of selection and an arbitrarily complex population size history. We then use this framework to show that the unconditional age distribution of negatively selected alleles is very closely approximated by re-weighting the neutral age distribution in terms of the negatively selected SFS, suggesting that allele ages provide very little information about the DFE beyond that already contained in the present day frequency. To confirm this prediction, we extended the standard Poisson Random Field (PRF) method to incorporate the joint distribution of frequency and age in estimating selection coefficients, and test its performance using simulations. We find that when the full SFS is observed and the true allele ages are known, including ages in the estimation provides only small increases in the accuracy of estimated selection coefficients. However, if only sites with frequencies above a certain threshold are observed, then the true ages can provide substantial information about the selection coefficients, especially when the selection coefficient is large. When ages are estimated from haplotype data using state-of-the-art tools, uncertainty about the age abrogates most of the additional information in the fully observed SFS case, while the neutral prior assumed in these tools when estimating ages induces a downward bias in the case of the thresholded SFS.

## 1 Introduction

Understanding how natural selection has shaped patterns of genetic variation is a core goal in population genetics (Wright, 1931; Fisher, 1930; Haldane, 1932). One valuable way of developing this understanding is by learning about a distribution of fitness effects (DFE), i.e., the probability distribution on the selection coefficients of newly arising mutations. (Eyre-Walker and Keightley, 2007; Boyko et al., 2008; McVicker et al., 2009; Coop, 2016; Brandvain and Wright, 2016; Gravel, 2016; Simons and Sella, 2016; Amorim et al., 2017; Booker et al., 2017; O’Connor et al., 2019; Durvasula and Lohmueller, 2021; Zeng et al., 2021; Simons et al., 2022). There are several approaches to estimating the DFE. For species amenable to laboratory experimentation, the DFE can be estimated directly by comparing the fitness of individuals carrying different mutation complements (Fowler et al., 2010; Hietpas et al., 2011; Bataillon and Bailey, 2014; Boucher et al., 2014), or indirectly from changes in the fitness of lines of laboratory organisms during mutation accumulation experiments (Eyre-Walker and Keightley, 2007; Halligan and Keightley, 2009). In many organisms (e.g., humans), this is not possible, and the DFE is estimated from polymorphism data using population genetic models (Bustamante et al., 2001a; Williamson et al., 2005; Kim et al., 2017; Johri et al., 2022).

These approaches typically use the site frequency spectrum (SFS)—a vector recording the number of mutations observed at each possible sample frequency—as a summary statistic for estimating the DFE under the Poisson random field (PRF) model first introduced by Sawyer and Hartl (1992). Intuitively, because negative selection against a mutation reduces the probability that it will climb to a high frequency in the population, and thus be observed at a high frequency in the sample, information about the DFE comes from the shape of the SFS. However, the frequency of a single variant provides relatively little information about its selection coefficient, requiring that information be pooled across a relatively large number of sites to estimate a DFE (Williamson et al., 2005; Boyko et al., 2008; Kim et al., 2017; Simons et al., 2022; Zeng et al., 2023). As a consequence, DFE inferences typically focus on relatively large classes of sites (e.g., all non-synonymous variants), and efforts to estimate the DFE for smaller subsets of sites (e.g., variants associated with a particular complex trait, Simons et al. 2022 or the set of loss-of-function mutations in a short gene, Zeng et al. 2023) come with significant uncertainty.

One plausible way of overcoming this limitation is to include more information about the past frequency trajectory of each variant into the inference. To this point, breakthroughs in coalescent inference methods over the past 10 years have enabled the estimation of gene trees from DNA sequence data (Rasmussen et al., 2014; Kelleher et al., 2019; Speidel et al., 2019; Albers and McVean, 2020; Lewanski et al., 2024), while the increase in the amount of ancient DNA sequenced over roughly the same time period has allowed for temporal sampling of allele frequencies. Recent work has leveraged both the estimated genealogies (Stern et al., 2019; Vaughn and Nielsen, 2023) and ancient DNA time series to estimate positive selection coefficients for individual beneficial mutations (Malaspinas et al., 2012; Mathieson and McVean, 2013; Irving-Pease et al., 2022; Mathieson and Terhorst, 2022).

By contrast, methods for incorporating such information into the DFE inference paradigm for sites under negative selection have largely done so implicitly, using custom-built models and/or simulations to link patterns of haplotypic variation to the DFE for deleterious mutations (Johri et al., 2020; Ortega-Del Vecchyo et al., 2022). Outside of the traditional DFE inference paradigm, one simple summary of an allele’s past trajectory, its estimated age, has been used explicitly to study the impact of negative selection on complex trait associated loci using GWAS data. In this context, allele age estimates have been used to develop annotations that explain spatial variation across the genome in the magnitude of the contribution to complex trait variance (Gazal et al., 2017, 2018; Hujoel et al., 2019; Kichaev et al., 2019; Kanai et al., 2021; Shi et al., 2021; Nait Saada et al., 2023). The sign of this association is consistently negative across traits, such that younger alleles contribute more to heritability, consistent with the impact of negative selection. Notably, this association is not explained by the mutual association of heritability and age with minor allele frequency, suggesting that the ages may contain substantial additional information about the impact of selection beyond that which is included in the allele frequency. However, we still lack a complete understanding of the relationship between frequency, age, and strength of selection, limiting our ability to clearly interpret these associations.

In this paper, we study the utility of including allele age into the inference of deleterious fitness effects. Allele ages represent an attractive choice for this purpose because 1) when combined with the present day frequency, they would seem to summarize much additional information about an allele’s past trajectory in a single data point, and 2) there has already been much theoretical and empirical work on the distribution of allele ages, which we can build on. For example, derived alleles under selection (whether positive or negative) are known to be younger on average than neutral alleles at the same frequency in the population (Kimura and Ohta, 1973; Maruyama, 1974; Kiezun et al., 2013, also see Figure S1). In fact, as famously shown by Maruyama (1974), after conditioning on the present day frequency of the allele in the population, the distribution of ages for alleles under positive and negative selection of the same magnitude are identical (assuming additivity), though this symmetry breaks down if we condition instead on sample frequency (Stephens and Donnelly, 2003), or if the population size is not constant over time (Ortega-Del Vecchyo et al., 2022).

Several authors have developed tools to approximate or simulate from the distribution of allele ages, either under neutrality (Rannala, 1997; Griffiths and Tavaré, 1998; Slatkin, 2000; Slatkin and Rannala, 2000), or in the presence of selection (Slatkin, 2001; Wiuf, 2001; Stephens and Donnelly, 2003). A general expression for the density on the age of selected alleles is given by Griffiths (2003) in terms of the fixation probability and transition density of the Wright-Fisher diffusion, though evaluating this expression requires an appropriate numerical approximation of the transition density (e.g., Song and Steinrücken, 2012). If the population size is constant through time, samples from the distribution of allele age conditional on the population frequency can be obtained by simulating allele frequency trajectories forward in time, starting from the desired frequency and recording the time until loss. Then, due to the time reversibility of the Wright-Fisher diffusion, the distribution of these times until loss represent valid samples from the distribution of allele ages (Slatkin, 2001). Crucially, when the population size is not constant, this time reversibility no longer holds and this method is not valid. In this case, samples from the age distribution can still be obtained by simulation, either via a brute force forward-in-time approach (using a tool like in Ortega-Del Vecchyo et al., 2016), or via an importance sampling approach introduced by Slatkin (2001). However, all of these simulation based approaches entail substantial computational cost and/or the potential for error due to the Monte Carlo approximation.

Here, we first build on the moments framework developed by Jouganous et al. (2017) to introduce an accurate and efficient numerical method for computing the distribution on the age of a segregating variant given its sample frequency, selection coefficient, and an arbitrarily complex population size history. Second, we use this framework to show that the age distribution of negatively selected alleles with a given scaled selection coefficient can be closely approximated by re-weighting the age distribution of neutral variants across allele frequency bins by the ratio of the normalized SFS entries for the deleterious and neutral variants. Notably, the same is not true for positively selected variants, where information about the past frequency trajectory of an allele has proven extremely valuable (Hejase et al., 2020). This observation suggests that, if the full distribution of allele frequencies is observed, then allele ages should carry relatively little information about negative selection coefficients beyond that which is already contained in allele frequency data, in contrast to the case for positively selected variants To verify this prediction, we extend the standard PRF model for estimating fitness effects from distributions of allele frequencies to include the joint distribution of frequency and age, using our numerical method. Finally, we use additional simulations to show how allele ages *do* provide useful information about negative selection coefficients when there is a threshold on the minor allele frequency (e.g., as in analyses of GWAS summary statistics).

## 2 Results

### 2.1 A numerical method for the density on allele ages

First, we develop a numerical method to compute the distribution on allele age for a segregating variant under selection, conditional on its sample frequency. To do this, we build on a numerical approximation to the Wright-Fisher diffusion developed by Jouganous et al. (2017) (see also Evans et al., 2007; Malaspinas et al., 2012, for closely related earlier work), which allows us to compute this density on age using an efficient dynamic programming algorithm.

To begin, we consider a large population of *N* diploid individuals under a model consistent with the Wright-Fisher diffusion with selection. We imagine tracking the evolution of a very large number of independent sites, each with a small mutation rate, such that an infinite sites approximation applies. At each site, we follow a sample of 2*n* lineages through time (assuming *n* ≪ *N*), and track the number of lineages at each site that carry a derived allele. Using the moment recursions developed by Jouganous et al. (2017), we can track the expected number of sites where the derived allele will be found *i* times in the sample, i.e., the expected site frequency spectrum. We write 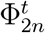 to denote the length 2*n* − 1 vector which records this expected SFS in generation *t*, where *t* counts down as time moves forward until *t* = 0 at the present generation. The *i*^*th*^ entry of 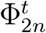 therefore gives the expected number of sites with a derived allele at frequency *i* in the sample of size 2*n, t* generations before the present. Given 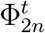, we can obtain the expected SFS in generation *t* − 1 via two steps. First, common ancestor events (i.e., genetic drift/coalescence) and selection events lead to the movement of mass among adjacent bins of the SFS (along with loss of mass from the singleton and “2*n* − 1”-ton bins) from one generation to the next. Second, new mutations are expected to arise at frequency *i* = 1 at 2*nµ*_*t*_ sites that previously lacked segregating derived alleles, where *µ*_*t*_ is the total mutation rate across all sites in generation *t*. These dynamics are described by

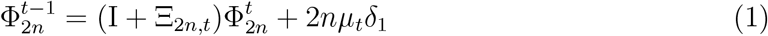

where I is the identity matrix, Ξ_2*n,t*_ is a tri-diagonal matrix containing the coefficients describing how the combination of genetic drift and selection operating in generation *t* move mass between adjacent bins in the expected site frequency spectrum, and *δ*_1_ denotes a vector with a 1 in the singleton bin and 0 elsewhere. The coefficients of Ξ_2*n,t*_ are precisely as given by Jouganous et al. (2017), including their jackknife approximation to close the moment equations in the presence of selection (we reproduce these coefficients in Section S2). Notably, differences in the population size or strength of selection across generations are accounted for via differences in the coefficients of Ξ_2*n,t*_.

The probability distribution on the allele age, *a*, given an arbitrary sequence of selection coefficients, population sizes, and mutation rates, can be written as

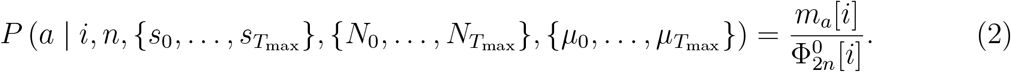

where *m*_*a*_[*i*] is the expected number of mutations that arise in generation *a* and are found at frequency *i* in the present day. We first compute the denominator, 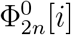, by initializing 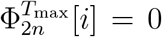 for all *i*, with *T*_max_ set sufficiently far in the past that a negligible number of mutations arising before this time would be expected survive to the present day. We then iterate Equation (1) until we obtain 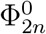. To compute the numerator, we imagine tracking the sample frequency of the the derived allele at each of the the 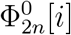 sites found at frequency *i* in the present, backward in time until for each one we encounter the mutation event from which it arose. Concretely, we write 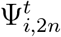 for the expected SFS of this conditional sample at time *t*, initializing

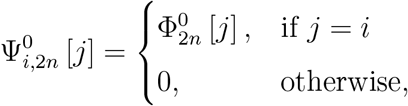

at generation 0. We then evolve this conditional SFS backward in time by iterating

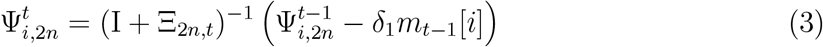

from *t* = 0 to *t* = *T*_max_, where

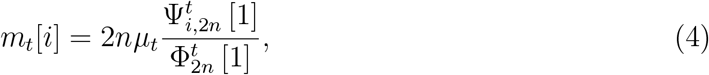

The ratio 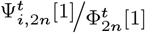 gives the expected fraction of mutations found as singletons in a sample taken in generation *t*, which are destined to be found at frequency *i* in the present, and can thus be interpreted as the probability that a random singleton at generation *t* will be found at frequency *i* in the present. The product of this probability with the number of mutations arising in generation *t* (i.e., 2*nµ*_*t*_) therefore gives expected number of mutations that arise in generation *t* and are found in the sample at frequency *i* in the present.

Notably, if the population size, selection coefficient and mutation rate are constant over time, then the 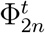[1] attains a steady-state value, Φ_2*n*_[1]. In this case, the *µ*_*t*_ are also the same for each generation, so Equation (2) reduces to

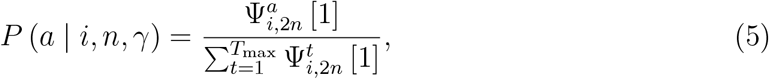

(where *γ* = 2*Ns* is the population scaled selection coefficient). This follows from the reversibility of the Wright-Fisher diffusion with respect to its steady state, and is related to the observation that the age distribution in a constant size population (with a constant selection coefficient and mutation rate) can be obtained by simulating forward in time until loss and then reversing the trajectory (Maruyama, 1974; Slatkin, 2001), while this is not possible in a population that varies in size, or if the selection coefficient or mutation rate vary over time. In principle, this suggests that if all three of these parameters are held constant, then the initial forward pass through time can be skipped, and that the age distribution can be obtained with only a single backward pass through time.

However, computing the 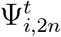[1] for each generation still requires that we compute the *m*_*t*−1_[*i*], so we still need to know the value of Φ_2*n*_ [1]. In principle, for constant size population under additive selection, Φ_2*n*_ [1] could be obtained by integrating the analytical expression for the population SFS (Wright, 1938a; Bustamante et al., 2001b) against the binomial probability of obtaining a singleton in the sample, but here we simply run the algorithm forward in time as in the non-equilibrium case, given that doing so is not computationally expensive.

This method provides a means to compute the density on allele age that is both accurate and efficient. For example, computing the distribution of ages for a single *i* in a sample of *n* = 125 and selection coefficient *s* = −5 *×* 10^−4^ assuming the piece-wise constant model of exponential growth inferred by Tennessen et al. (2012) for African-American individuals (over 55,000 generations, shown in Figure S2) takes on average 4 seconds on a MacBook Pro M1 (2021). In Figure S3, we validated the accuracy of the method by comparing the cumulative distribution of ages conditional on segregation in a sample of *n* = 125 to that obtained via forward-in-time simulations using `PReFeRSim` (Ortega-Del Vecchyo et al., 2016) under two scenarios: neutrality (*s* = 0) in a population of constant size, and moderate negative selection (*s* = −5 *×* 10^−4^) in the Tennessen et al. (2012) growth model. In both cases, the distribution obtained with our method closely matches the simulations (see Figure S3). Notably, although the method is fast to obtain the age distribution for a single sample frequency, obtaining age distributions for all sample frequencies, takes 2*n* times as long, because the age distribution must be computed separately for each sample frequency. This can become substantial, especially if sample sizes are large (e.g., ∼ 1,000 seconds for the example case with *n* = 125 considered here). In Section S1, we outline an alternative algorithm to obtain the age distribution for all sample frequencies at the same time in a more efficient manner. Briefly, this method relies on the fact that the probability that an allele found at frequency *i* is *a* generations old is proportional to the number of such mutations present in the sample. Thus, we can efficiently compute the density on allele age across all frequencies by first computing the sample SFS conditional on allele age, forward in time, for each generation in which a mutation could have arisen. For the example case considered here, this method takes approximately 30 seconds to obtain all 2*n* − 1 = 249 age distributions. Both methods scale linearly in sample size and *T*_max_ and the runtimes for different combinations of parameters are shown in Figure S4.

### 2.2 The age distribution of selected alleles

We next applied our method to study how selection impacts the distribution on ages, conditional on segregation in a present day sample. In this section, we focus largely on the constant size case, so we represent time in coalescent units and the strength of selection in terms of the population scaled selection coefficient, *γ* = 2*Ns*, and we write the density on age given a particular scaled selection coefficient as *P* (*a* | *i, n, γ*). In Figure 1, we plot age distributions in a sample of size *n* = 125, across a range of selection coefficients and sample frequencies. As expected, for a given sample frequency, alleles under stronger selection are younger on average with less variation in age than less strongly selected alleles (Maruyama, 1974; Wiuf, 2001; Griffiths, 2003; Stephens and Donnelly, 2003, also see Figure S1).

**Figure 1:**
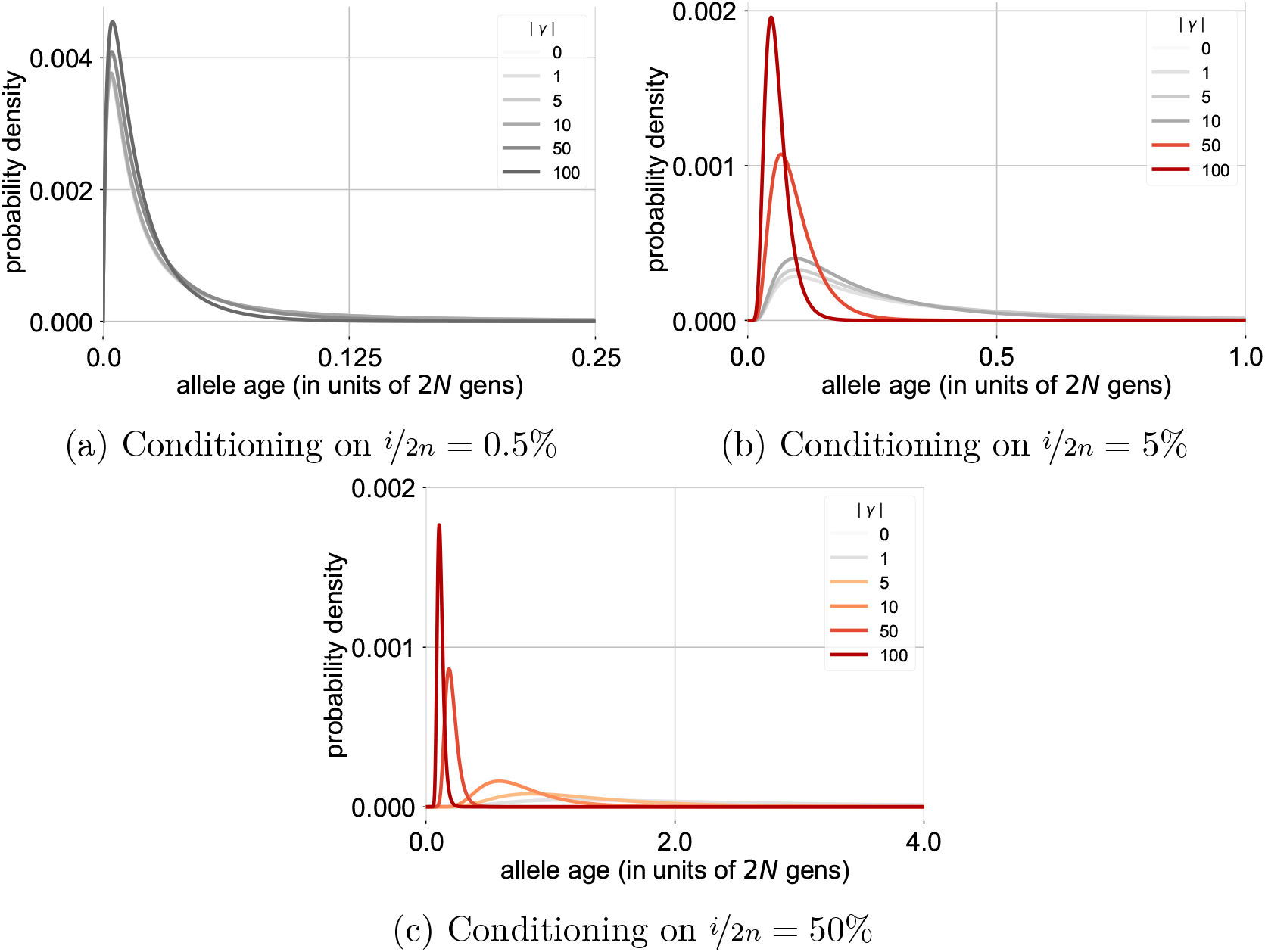
Age distributions under different strengths of selection, conditional on segregating at a particular frequency in a diploid sample of *n* = 250. For a particular sample frequency *x*^⋆^, scaled selection coefficients less than 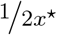 are shown in gray-scale, while the selection coefficients that are larger than this threshold are shown in color (in an orange-red scale).

More precisely, for a given sample frequency, the allele age distribution for sites under weaker selection is generally very similar to the age distribution for neutral alleles with the same sample frequency, with significant deviations arising only when selection is stronger. This pattern is consistent with the impact of selection on the SFS. Specifically, to a first approximation, the population SFS for sites with scaled selection coefficient *γ* resembles the neutral SFS at frequencies below a threshold value of 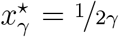, but drops off quickly above this value (Wright, 1938b, also see Figure S5). The sample SFS is expected to show a similar pattern, so long as 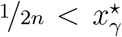, such that sites with population frequencies both above and below this threshold are represented in the sample. Thus, we would expect that 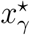 should also mark the approximate boundary between sample frequencies for which the age distribution is similar to that of neutral alleles, and those for which it is not. To test this prediction quantitatively, we use our numerical framework to compute the Kullback-Leibler (KL) divergence between the conditional age distribution of selected alleles at a given sample frequency and the corresponding age distribution of neutral alleles at the same sample frequency,

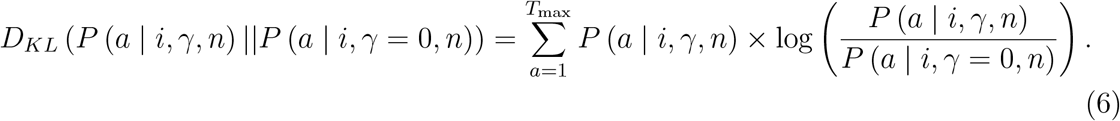

In Figure 2, we plot this divergence as a function of the scaled selection coefficient, *γ*, and sample count, *i*, assuming that *n* = 100. Confirming our prediction, we find that broadly, across the range of scaled selection coefficients, the divergence is close to zero for sample frequencies less than 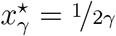 (shown by the gray dashed line), but begins to increase above this value. This observation suggested that it should be possible to closely approximate the unconditional age distribution of negatively selected alleles by a simple re-weighting of the neutral age distribution in terms of the SFS for negatively selected alleles. Specifically, we hypothesized that *P* (*a* | *γ, n*), the unconditional density on allele age given the scaled selection coefficient, can be well-approximated as

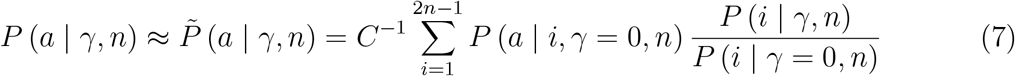

where *P* (*i* | *γ, n*) and *P* (*i* | *γ* = 0, *n*) are the expected sample SFSs for selected and neutral alleles respectively (normalized to probability mass functions conditional on segregation), *P* (*a* | *i, γ* = 0, *n*) gives the age distribution of neutral alleles at sample frequency *i*, and 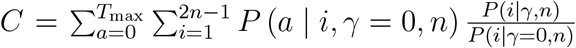 is a normalizing constant which ensures that 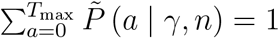.

**Figure 2:**
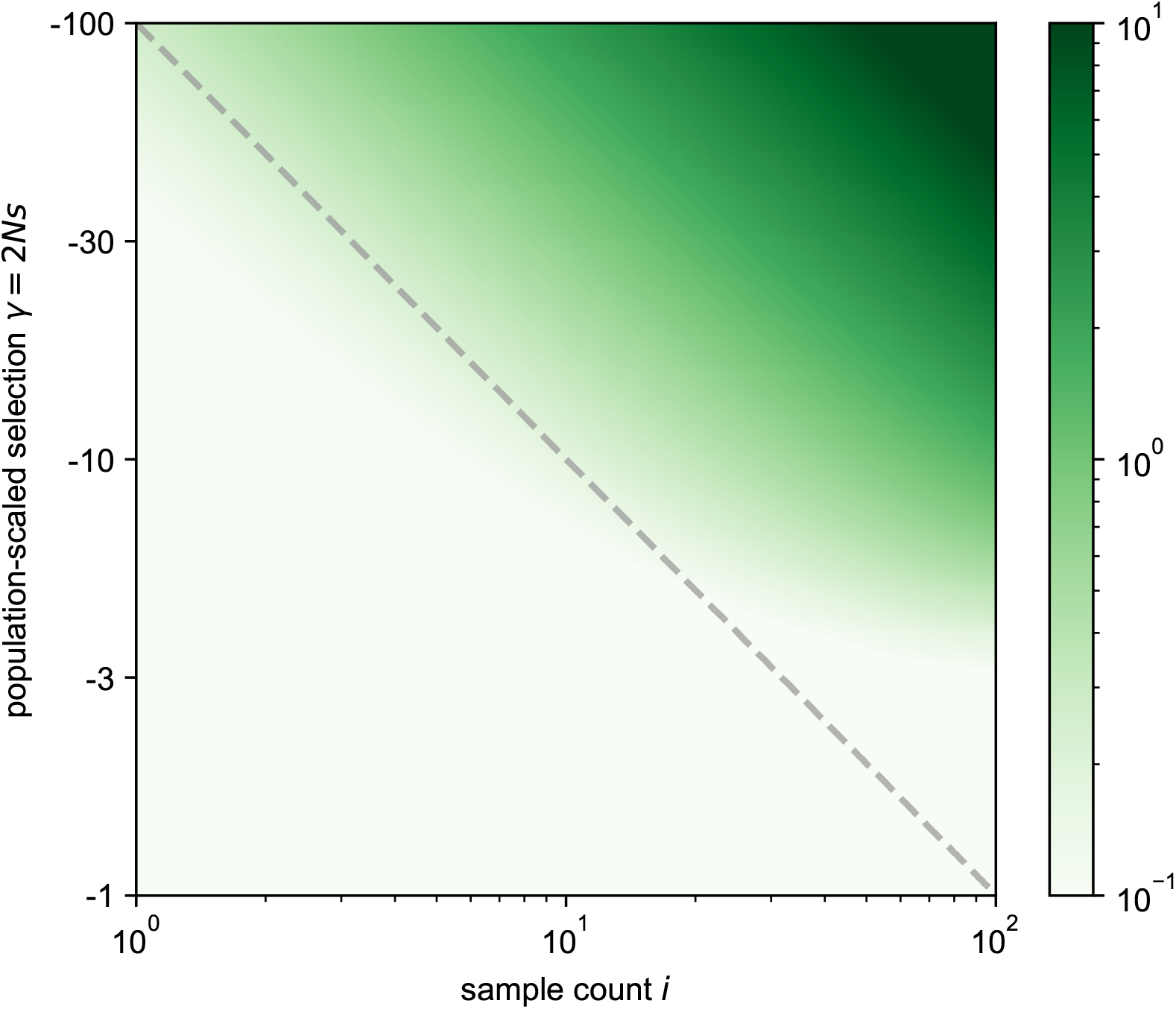
The heatmap of Kullback-Leibler (KL) divergence between the age density of alleles given a particular selection coefficient, *P* (*a* | *i, γ, n*), and that of neutral alleles, *P* (*a* |*i, γ* = 0, *n*), conditional on the sample allele count *i*. The gray dashed line indicates the threshold, 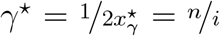, above which we expect the conditional age distribution of selected alleles to differ substantially from that of neutral alleles. Values in this figure are calculated for a diploid sample size of *n* = 100, so a sample count *i* = 100 corresponds to a sample frequency of 50%.

In Figure 3, we compare this approximation to the exact unconditional age distribution, which we compute using our numerical framework as

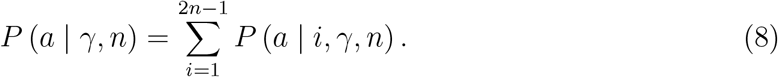

Figure 3a shows this comparison for a representative case of moderate negative selection (*γ* = −20). In this case, the approximation in Equation (7) closely matches the truth in Equation (8), with significant differences emerging only for older alleles which are unlikely to be found under either the true model or the neutral resampling approximation (see Figure S6 for cases with *γ* = {−100, −10, 0, 10, 100}). This pattern indicates that the difference between the unconditional age distribution for neutral alleles and those under moderate negative selection is mostly explained by the relative dearth of high frequency alleles under negative selection, and not by the differences in the conditional age distributions.

**Figure 3:**
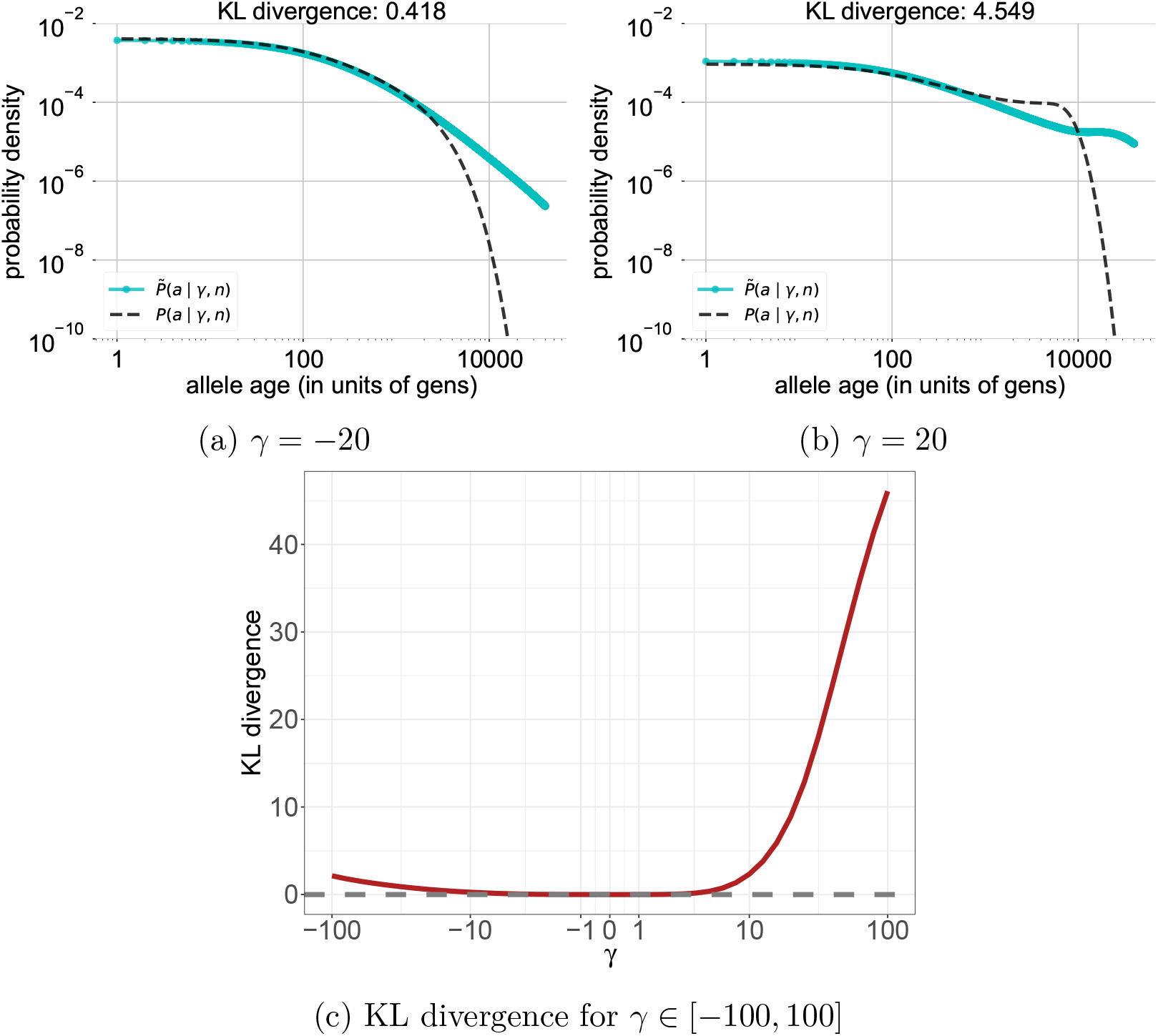
The KL divergence between the unconditional age distributions of a particular selection coefficient (Equation (8)) and the age distribution approximated by downsampling from the neutral frequency spectrum (Equation (7)) across a range of selection coefficients. In **a)**, we observe that for moderate negative selection (*γ* = −20), the two distributions are very similar for young alleles and differ only for the oldest alleles of which there are very few. In **b)**, for sites experiencing moderate positive selection (*γ* = 20) the agreement between the true and approximated age distribution is much worse than for negative selection. In **c)**, we plot the KL divergence between the true and approximated age distribution (Equation (6)) across a range of selection coefficients.

In contrast, for a complementary case of moderate positive selection (*γ* = 20; Figure 3b), Equation (7) provides a poor approximation of the true unconditional age distribution. The reason is that positive selection amplifies the abundance of precisely those frequency bins for which the conditional age distributions are most strongly shifted relative to the neutral expectation. As a result, the neutral resampling distribution places greater mass on older ages compared to the true distribution, given the abundance of higher frequency alleles in this positively selected case compared to the previous negatively selected case.

To quantify the accuracy of this approximation, we again compute a KL divergence, this time between the approximation given in Equation (7) and the exact unconditional age distribution in Equation (8). The relevant KL divergence is then given by

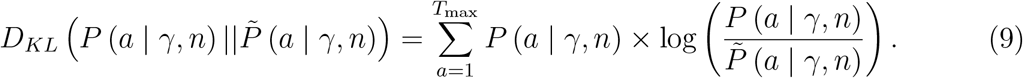

In Figure 3c, we plot the divergence as a function of the scaled selection coefficient for *γ* ranging from −100 to 100. In general, the divergence is very low for negatively selected alleles, and begins to increase substantially above zero only once selection becomes quite strong (e.g., *γ <* −50). In contrast, the divergence increases rapidly with the strength of positive selection, consistent with our expectation.

### 2.3 Utility of allele ages in DFE inference

We now turn our focus to directly assessing the utility of allele age estimates for estimating the strength of negative selection acting on a variant or set of variants. Our results in the previous section show that for negatively selected alleles, once we condition on the distribution of allele frequencies, the distribution of ages is relatively insensitive to the precise value of the selection coefficient. This suggest that once the frequency of a variant under negative selection is measured, its age is unlikely to contain much additional information about its selection coefficient. To test this prediction, we used our numerical framework to implement simple inference frameworks for estimating selection coefficients using **1)** only the site frequency spectrum (SFS), or **2)** the joint distribution of site frequencies and ages. Both implementations use the standard Poisson Random Field (PRF) model introduced by Sawyer and Hartl (1992) to compute the likelihood of the selection coefficient (see Methods).

To compare the two approaches, we performed an experiment in which we simulated paired allele frequencies and ages for 1,000 unlinked sites under additive selection using `PReFerSim` (Ortega-Del Vecchyo et al., 2016) across a grid of scaled selection coefficients ranging from *γ* = *−*100 to *γ* = 100. We then inferred the value of *γ* via maximum likelihood using both frequency only and frequency & age. For each value of *γ*, we replicated this procedure 100 times, and visualized the distribution of MLEs across these 100 replicates in Figure 4. The estimates from both approaches are largely unbiased, with the exception of a slight downward bias for the frequency only approach in the case of weak positive selection. To determine how much additional information is contained in the ages, we compared the variance of the MLEs across replicates (i.e., the squared standard error) using the two different approaches. We find that even when the allele ages are known exactly, including them in the inference results in only a small decrease in the standard error relative to the frequency-only approach for negatively selected alleles, consistent with our predictions above, and in contrast to positively selected alleles, where ages provide a significant benefit (Figure 4c). We observed a similar pattern when both the simulations and the inference are performed under the Tennessen et al. (2012) piece-wise constant model of exponential growth (Figures S7 & S8).

**Figure 4:**
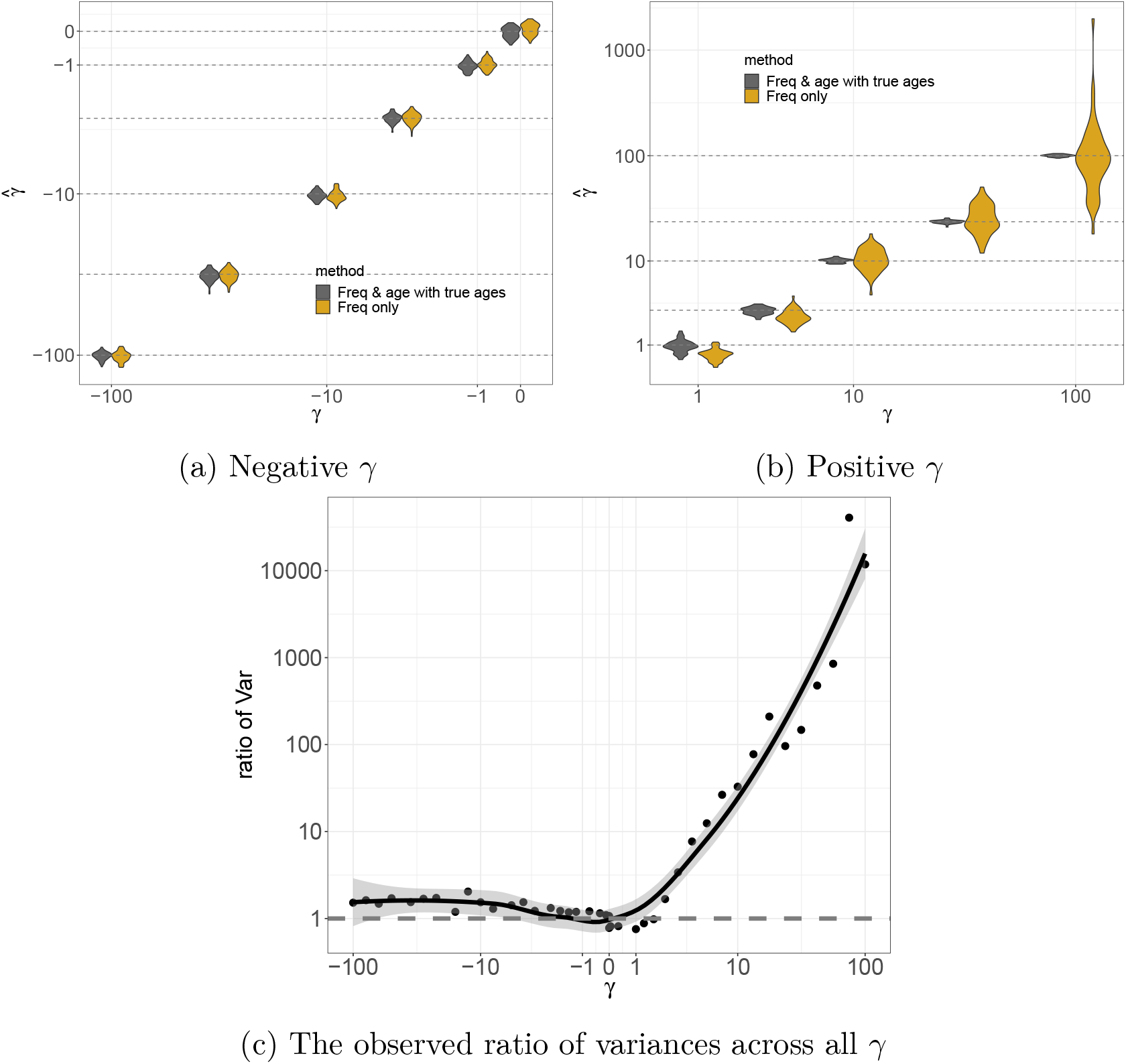
Selection coefficient estimation for a constant demographic history of *N* = 10,000 using data simulated with `PReFerSim` (Ortega-Del Vecchyo et al., 2016). **a), b)** Violin plots showing accuracy of estimation for different values of population-scaled selection coefficient *γ* using allele frequency & age data versus allele frequency alone. The X-axis shows different values of simulated *γ*, while the Y-axis shows the distribution on estimated 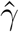 over 100 independent replicates. The dashed horizontal lines denote the simulated values to aid in visualizing bias. On the negative side of the spectrum, we found that the MLE are close to the true value in both cases, with the approach including ages having slightly smaller error bars indicating more information about the selection coefficient in the data (especially for stronger values of selection). **c)** Loess line fitting the ratio of variance (squared standard error) estimates calculated using Equation (15) from the frequency only approach and the frequency & age approach, for *γ* ∈ [−100, 100].

In Figure 3, we showed that age distributions conditioned on sample frequency differ significantly from neutral expectations only for variants above a threshold frequency that depends on the scaled selection coefficient. Given this observation, we hypothesized that allele ages would be of greater benefit in samples with a threshold on the SFS (as is often the case for genome-wide significant GWAS associations). We tested this prediction by inferring selection coefficients from simulated data with a MAF cutoff of *x*^⋆^ = 0.025 in a sample of *n* = 100. For scaled selection coefficients greater than one over twice the frequency threshold (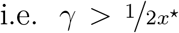) including ages in the inferences results in a sizeable increase in accuracy, but had little effect for scaled selection coefficients selection below this threshold (Figure 5b).

**Figure 5:**
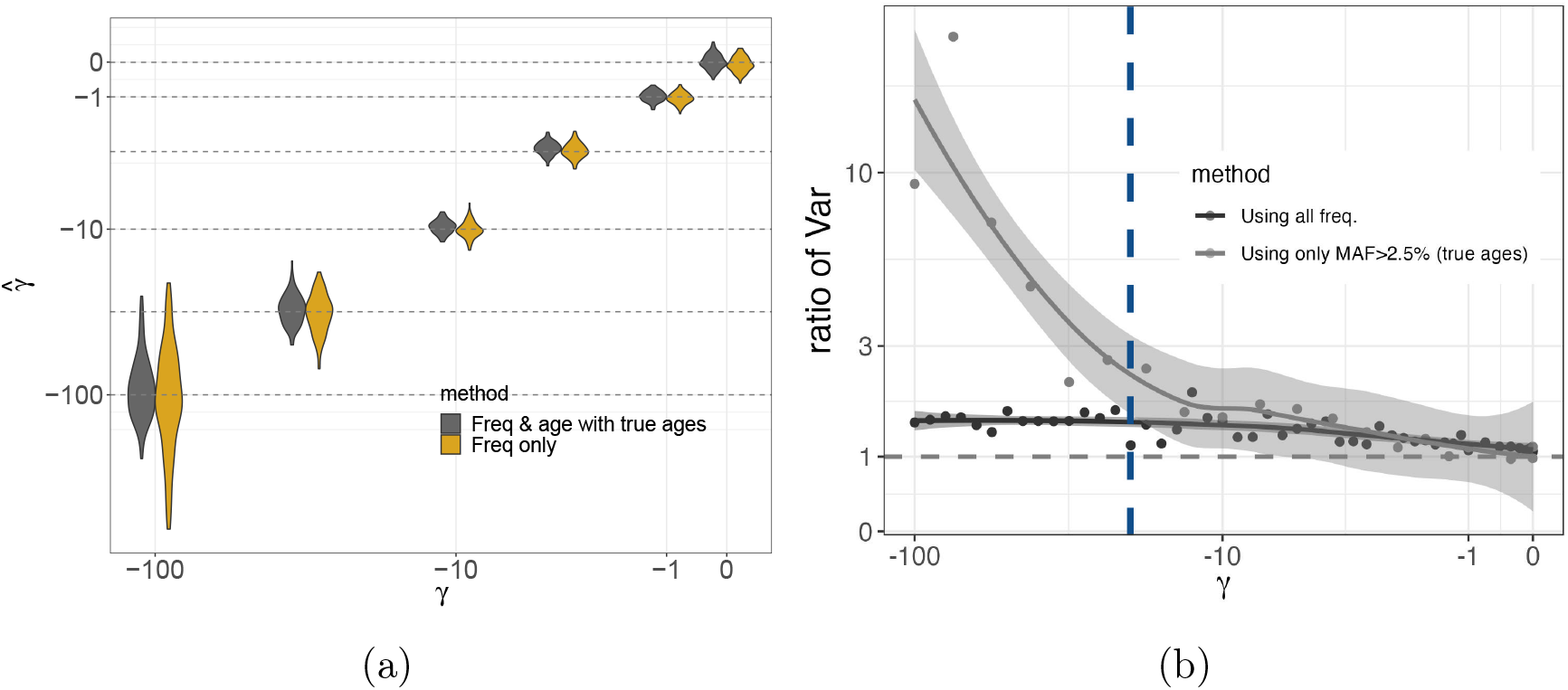
Selection coefficient estimation for neutrality and negative selection for a constant demographic history of *N* = 10,000 using data simulated with `PReFerSim` (Ortega-Del Vecchyo et al., 2016), using only sites with MAF≥2.5%. **a)** Violin plots showing the distribution of estimates across replicates when using the thresholded SFS, with or without ages. Estimates from both approaches are similarly unbiased across the entire tested range. **b)** Ratio of variances and a loess fit (similar to Figure 4c) to illustrate the gain in information due to including ages when there is a threshold on the SFS. We observe a significant increase in information gain for |*γ*| > ^1^*/*_(2*×*0.025)_ = 20 (indicated by the dashed blue line).

In practice, ages are typically also estimated from haplotypic data under a neutral prior. Conditional on the inferred tree, the posterior density on a mutation’s age then is uniformly distributed along the branch on which it arises. This induces an upward bias for selected alleles (Maruyama, 1974; Kiezun et al., 2013). To understand how this would impact inference of negative selection coefficients, we used `mssel` (Hudson, 2002) to simulate haplotypic data consistent with negative selection acting at a focal site. We then used `Relate` (Speidel et al., 2019) to estimate a single gene tree for each simulated focal site, and then used the allele ages implied by these trees to estimate selection coefficients. In the simplest method, we used as a point estimate the midpoint of the branch containing the focal mutation. This estimate is the posterior mean under a neutral prior, conditional on the estimated tree, and unsurprisingly results in selection coefficient estimates that are biased toward zero (Figure 6a). To better account for uncertainty in the estimated age, we next considered averaging over the uniform (neutral) posterior density on age on the branch on which the mutation arose. Interestingly, this approach was sufficient to remove the bias in the selection coefficient estimates, even though the age estimates still rely on a neutral prior (see Figure 6a). However, the added uncertainty about when the mutation arose is sufficient to abrogate what little improvement in accuracy the ages had provided when the full SFS is observed (Figure 6b).

**Figure 6:**
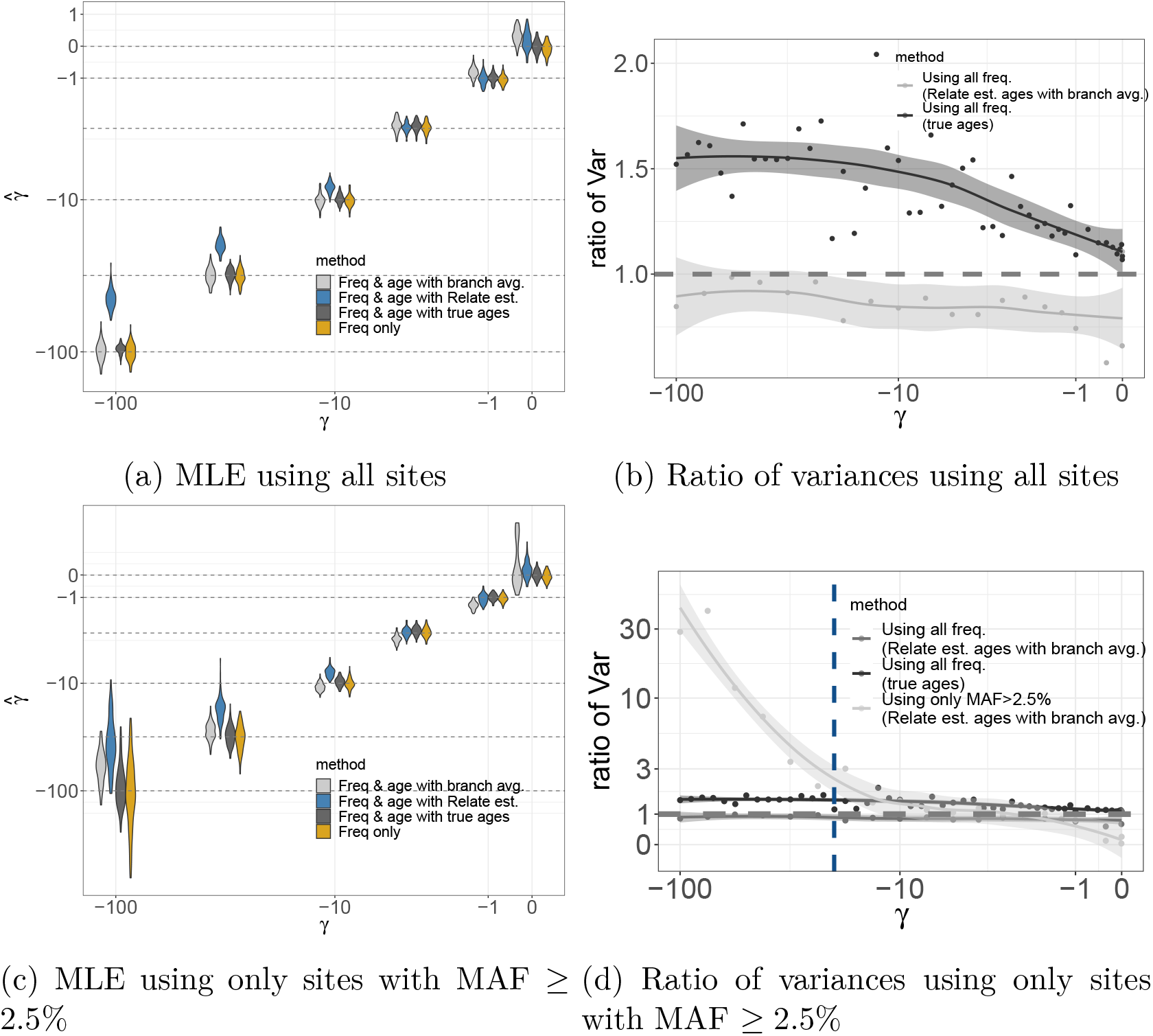
Selection coefficient estimation accuracy for neutrality and negative selection under a constant demographic history with *N* = 10,000 using data simulated using `PReFerSim` (Ortega-Del Vecchyo et al., 2016) and haplotypes simulated using `mssel` (Hudson, 2002) and ages estimated using `Relate` (Speidel et al., 2019). **a)** Using sites at all frequencies and raw age estimates from `Relate`, estimated selection coefficient are biased toward zero due to the neutral coalescent prior. However, this bias is eliminated by averaging over the density on age on the branch which it arose versus using the point estimate from `Relate`. **b)** However, uncertainty in the age estimates (i.e., increase in length of estimated branches) also erases nearly all of the additional information gained by including age estimates. **c)** When using the SFS thresholded at MAF ≥ 2.5%, estimates using the age density averages across the estimated branch also become biased, especially for larger scaled selection coefficients. **d)** Despite the bias seen in panel **c**, including ages still substantially reduces the variance of the estimates when the SFS is thresholded and ages are estimated using `Relate`.

Consistent with our observations for true ages, when there is a threshold on the SFS such that only sample frequencies greater than *x*^⋆^ = 0.025 are observed, *and* the ages are estimated, then using the ages does result in a substantial decrease in the standard error of the estimated selection coefficients (Figure 6c). However, in this case, averaging the inference over a uniform distribution on allele ages given the inferred tree does not fully remove the downward bias in the estimated selection coefficients, particularly when the scaled selection coefficient is large relative to the frequency threshold (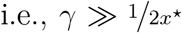; Figure 6d). Intuitively, this is because the sites included in the inference, which all have frequencies just above the threshold frequency *x*^⋆^, are precisely the ones for which the true trees deviate most significantly from the expectation under the neutral prior. Fully accounting for this effect would require an importance sampling scheme that averages over trees while upweighting contributions from those with younger allele ages (Coop and Griffiths, 2004; Stern et al., 2019; Vaughn and Nielsen, 2023). Such an approach could be implemented in our sample based scheme (see Discussion), but we do not pursue it here.

## 3 Methods

### 3.1 PRF Model

The standard PRF model for the SFS as introduced in Sawyer and Hartl (1992) is given by

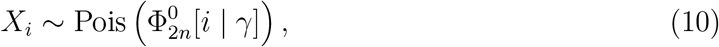

where *X*_*i*_ is the number of sites observed with sample frequency *i* and 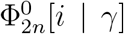 is the expected number of sites at sample frequency *i* in the present generation given the scaled selection coefficient *γ*. We note that 1) elsewhere in the manuscript, we suppress this conditioning in our notation, writing just 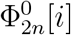, but here we make it explicit so that its role in the inference is clear, and 2) although we write this section in terms of the *constant size* (equilibrium) case for simplicity, the extension to the non-equilibrium case is straightforward.

This framework can be expanded to the joint distribution of frequency and age via a simple Poisson splitting argument. If we write *X*_*ia*_ for the number of sites observed to have frequency *i* and age *a*, then the distribution of *X*_*ia*_ arises from splitting the *X*_*i*_ across age bins, so that

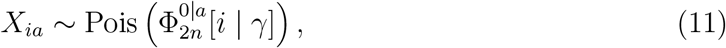

where 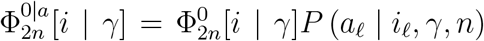 is the expected number of such sites given a selection coefficient *γ* (see also Section S1, where we provide a method to compute 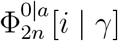 directly). The likelihood of the selection coefficient can thus be written

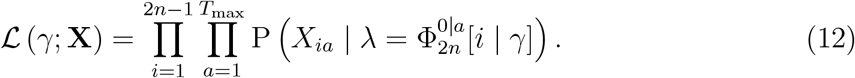

where 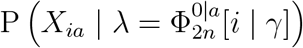 is the Poisson probability of *X*_*ia*_ given 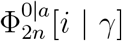. Alternatively, we can also rewrite (12) explicitly in terms of the Poisson splitting argument for the *X*_*i*_, as

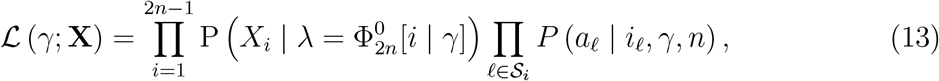

where 𝒮_i_ indicates the set of sites found at frequency *i* in the sample.

In practice, existing gene tree estimation methods (e.g., `Relate` from Speidel et al. (2019), `GEVA` from Albers and McVean (2020), `tsdate` from Wohns et al. (2022), etc.) employ a neutral coalescent prior, and thus provide biased age estimates for alleles under selection. To counteract this effect, we apply a branch averaging scheme which integrates over the probability of observing an allele with a particular frequency and age given by the duration of the entire branch on which it arose.

Concretely, for a given site 𝓁 (and a tree estimated via the corresponding haplotypes), we draw *D* mutations (each indexed by *d*) uniformly across the branch on which the allele arose

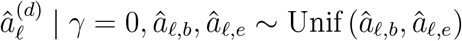

where *â*_𝓁,*b*_ is the most recent generation in which the branch exists, and *â*_𝓁,*e*_ the most ancient generation in which it exists.

Now, our likelihood for the selection coefficient includes an integration over the density on age given a selection coefficient and a branch (i.e., tree),

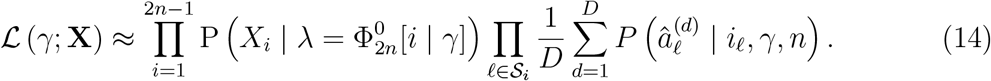

This equation differs from Equation (13) in that it simply replaces the probability of observing a single value with a distribution over a set of values.

### 3.2 Simulation and estimation framework

The goal of our simulations was to produce paired data of allele frequency & age {*i*_𝓁_, *a*_𝓁_} for a site 𝓁 given a population-scaled selection coefficient *γ* = 2*Ns* (assuming additive selection, 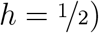 and demographic history {*N* }. We used the forward-in-time simulator `PReFerSim` to generate unlinked sites under each scenario due to its speed and its ability to record ages and trajectories of selected alleles (Ortega-Del Vecchyo et al., 2016). We tested the model under two demographic scenarios: a *constant size* case, where we set *N* = 10,000 for 100,000 generations (sufficient to reach approximate equilibrium for the entire range of simulated selection coefficients), and a *piece-wise exponential growth* case inferred in Tennessen et al. (2012) for humans over the last 50,000 generations (‘Africa 1T12’ from Adrion et al., 2020, see Figure S2). To capture the entire range of selection strengths, we simulated under 30 different selection coefficients, *s*, equally spaced on a log-scale from 5 *×* 10^−7^ (very weak, corresponds to *γ* = 0.01 in the *constant size* case) to 5 *×* 10^−3^ (strong, corresponds to *γ* = 100 in the *constant size* case), and, *s* = 0. This was repeated on both sides of the spectrum. The population scaled mutation rate *θ* was varied from 400 for *s* = *−*5 *×* 10^−3^ to 10 for *s* = 5 *×* 10^−3^, so as to obtain a similar number of segregating sites (≈ 1,000) across all selection coefficients. This was to ensure that the information about the selection coefficient came from the frequency and age, and not from the total number of segregating sites in the data.

Secondly, to explore the performance of the model in the case of biased (or estimated) ages, we used `PReFerSim` to record entire trajectories of alleles, which was used as input to the program `mssel` (Hudson, 2002) that outputs haplotypes containing the selected allele. These haplotypes were passed into `Relate` (Speidel et al., 2019) to construct gene trees at the selected sites and subsequently *estimate* ages of the selected mutations. To understand the extent of the bias under these estimated ages, we used the midpoint of the corresponding branch in the selection coefficient estimation framework instead of the true ages from before. To mitigate this bias for alleles under negative selection, we used the likelihood in Equation (14) with *D* = 500 mutations uniformly on the branch on which the mutation arose, as we found that the estimate of the selection coefficient did not change significantly with a higher number of draws. This exploration was done only for the *constant size* case, as we expect the findings to extend to more complex demographic histories.

For each combination of selection coefficient and population size history, we ran 100 replicates to get an accurate measure of the first two moments (mean and variance) of the distribution of the estimated selection coefficient, *ŝ*. In all cases, we used a maximum likelihood based approach to estimate the selection coefficients. Since we were optimizing over a single dimension, we used Brent’s method (default) in `scipy` (Virtanen et al., 2020) to minimize the negative log-likelihood of the data under the appropriate models. This could be extended to estimate multiple selection parameters (for instance, shape and scale of a gamma DFE).

#### 3.2.1 Measuring ratio of variances

The amount of information contained about a parameter (here, *γ* = 2*Ns*) in a certain type of data (say, SFS) is proportional to the width of the standard errors around the MLE–the lower the variance of the estimate around the MLE, the more the information about the parameter in the data (Lehmann and Casella, 2006). To measure the gain in information on adding allele ages, we compute the ratio of the empirical estimate of the two variances around the MLE (*ŝ*^*f*^ for frequency only and *ŝ*^*a*^ for frequency & age) calculated using the appropriate likelihoods from Equations (10) & (11), respectively (and where appropriate, Equations (13) or (14)). The higher the ratio of squared standard errors, the more the information that we have about the parameter of interest, with a value of 1 indicating equal amounts of information about the parameter in the two data sets. This ratio can be computed for each selection coefficient in our pre-specified range. Here, *j* reflects the index of the simulation replicate of the 100 replicates in total,

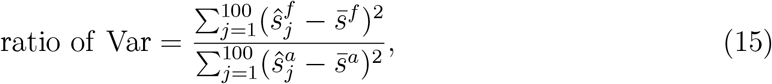

where 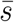 is the mean MLE across all replicates.

## 4 Discussion

Here we develop efficient numerical methods for computing the distribution of ages for selected alleles. This method improves on prior approaches to this problem by avoiding the computational cost and Monte Carlo error associated with simulations, producing accurate numerical approximations in seconds. Our work also builds on previous methods which have aimed to use the information about allele ages contained within patterns of haplotype variation to learn about the distribution of selection coefficients acting on negatively selected alleles (Kiezun et al., 2013; Johri et al., 2020; Ortega-Del Vecchyo et al., 2022). Interestingly, we find that if all bins of the SFS are observed in a reasonably large sample, then allele ages provide relatively little additional information about negative selection coefficients, particularly when we account for the fact that ages are estimated with error. However, incorporating ages can provide larger benefits if the frequency spectrum is truncated, e.g., in statistical genetics analyses of “common variants” (e.g., Gazal et al., 2017; Kichaev et al., 2019; Simons et al., 2022). Notably, because our method separates the problem of inferring allele ages from the problem of learning about selection coefficients conditional on the inferred allele ages, our results capture fundamental limits which cannot be overcome via improvements to the methods for inferring ages.

Nonetheless, there are several ways that our methods could be extended or improved upon. For example, although we focus on models with only a single population, the `moments` framework can accommodate multiple populations related via ancestral population splits, admixture events, and continuous migration. Computing the distribution of allele ages conditional on the sample frequencies observed across multiple populations should therefore be relatively straightforward, merely requiring additional bookkeeping to account for which population the allele ultimate arose in. Another plausible extension would involve replacing the Wright-Fisher diffusion by the discrete time Wright-Fisher model using methods recently developed by Spence et al. (2023). This approach could be used either to obtain the distribution on age given the population frequency, or by sub-sampling from this population SFS, the sample frequency that we focus on here. This approach would have the benefit that it would not rely on the assumption inherent to the Wright-Fisher diffusion that there is at most one coalescent event in the history of the sample per generation, making it more amenable to stronger selection and very large samples, at the cost of the increased computational expense required to track the full population and to compute a broader transition distribution in each generation.

Another way of improving upon our method would be to incorporate additional information about the frequency trajectory beyond that contained in just the age of the allele. Several such methods have been developed in the context of both temporal sampling/ancient DNA (Bollback et al., 2008; Malaspinas et al., 2012; Mathieson and McVean, 2013; Irving-Pease et al., 2022; Mathieson and Terhorst, 2022), or coalescent inference (Stern et al., 2019), or both (Vaughn and Nielsen, 2023). We expect that it would be possible to extend our method in either of these directions. The possibility of extending our method to full coalescent inference is particularly interesting. Specifically, the expressions required for sampling from the ancestral selection graph (ASG, Neuhauser and Krone, 1997) conditional on the present day sample configuration in a population at demographic equilibrium depend on ratios of the stationary sampling probabilities of the Wright-Fisher diffusion (Stephens and Donnelly, 2003). Despite the elegance of this approach, the need to simulate many lineages in the ASG which are not ancestral to the sample makes it computationally burdensome relative to the state-of-the-art structured coalescent HMM method (Stern et al., 2019; Vaughn and Nielsen, 2023), which explicitly models the latent population allele frequency. The requirement that these “virtual” lineages be simulated to obtain a valid sample stems from the fact that the outcome of a given selection event depends on the current frequency of the allele in the population, and thus cannot be determined until the outcomes of mutation and selection events occurring at earlier time points are known (Slade, 2000). However, in preliminary work in this direction, we have found that if the stationary sampling probabilities can be replaced with time-varying sampling probabilities which have already accounted for the distribution of times at which the mutation could have arisen (i.e., the distribution of allele ages), then the need to simulate virtual lineages can be avoided. Alternatively, it may be possible to compute the probability of a genealogy under selection via a purely forward-in-time approach, using a combination of the Moran model framework implemented in `momi` (Kamm et al., 2017) and the jackknife approximated selection operator introduced by Jouganous et al. (2017). Whether either of these approaches would offer benefits relative to the structured coalescent HMM method is an interesting question for future work.

However, given our focus on negative selection in this paper, it is worth asking whether additional information about the frequency trajectory beyond that already contained in the present day frequency and the age (e.g., frequencies at intermediate time points), to contain much additional information about the strength of negative selection experienced by an allele. Conditional on the end-points of a frequency trajectory, a larger selection coefficient means that a larger fraction of the frequency change required to go from one end-point to the other occurs closer to the present (Schraiber et al., 2013). This, in turn, would lead to coalescent events on the derived background being shifted closer to the present. However, a larger selection coefficient also means that segregating deleterious alleles will be younger on average (Kimura and Ohta, 1973; Maruyama, 1974; Kiezun et al., 2013), so there is less time for selection to substantially alter the frequency trajectories. Put differently, conditional on segregation, most deleterious alleles exist at frequencies below the 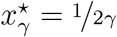 threshold at which the frequency spectrum transitions away from neutral behavior, so their trajectories should look approximately neutral. We would therefore expect to gain little additional information from their genealogies beyond what is contained in the two end-points. However, similar to the case with the ages, if we restricted the sample to sites with frequencies above this threshold, then we might expect the genealogies, or other information about the frequency trajectory, to be more useful.

Another setting where our approach may have greater utility is in the inference of selection on transpoable elements (TEs). For example, using neutral coalescent theory (Blumenstiel et al., 2014) and simulations (Horvath et al., 2022), prior work has shown that using an “age-adjusted” SFS (essentially, a binned version of the age conditioned SFS that we consider in Section S1) could be beneficial for estimating selection coefficients in TE families where the impact of selection is confounded by time-inhomogenous bursts of replication. Although we do not explore this direction in this paper, time-varying rates can be readily incorporated into our framework. More generally, our work illustrates how the now well-established framework of recursions for the SFS can be leveraged to address fundamental questions in population genetic inference.

## 5 Data Accessibility

All data utilized in this study were generated through computational simulations. The simulation scripts and source code are available in this repository: https://github.com/VivaswatS/selCoefEst.git. The source code for the program `mssel` is available at https://github.com/dortegadelv/HaplotypeDFEStandingVariation/tree/master/Programs/Mssel.

## 6 Author Contributions

VS and JJB contributed equally to the manuscript.

## 7 Acknowledgements

We would like to thank members of the Berg, Novembre, and Steinrücken labs, as well as members of the University of Chicago Program in Computational Biology (PCB) community for helpful discussions and feedback during the development of this project. We thank Aaron Ragsdale and Diego Ortega-Del Vecchyo for help with implementation and simulation. Additionally, we thank John Novembre, Matthias Steinrücken, and Xuanyao Liu for support at all stages of this work, and Carl Veller for comments on the manuscript. Computing was performed on servers maintained by the University of Chicago Research Computing Center.

## References

Adrion, J. R., C. B. Cole, N. Dukler, J. G. Galloway, A. L. Gladstein, G. Gower, C. C. Kyriazis, A. P. Ragsdale, G. Tsambos, F. Baumdicker, J. Carlson, R. A. Cartwright, A. Durvasula, I. Gronau, B. Y. Kim, P. McKenzie, P. W. Messer, E. Noskova, D. Ortega-Del Vecchyo, F. Racimo, T. J. Struck, S. Gravel, R. N. Gutenkunst, K. E. Lohmueller, P. L. Ralph, D. R. Schrider, A. Siepel, J. Kelleher, and A. D. Kern (2020, jun). A community-maintained standard library of population genetic models. eLife 9, e54967.

Albers, P. K. and G. McVean (2020). Dating genomic variants and shared ancestry in population-scale sequencing data. PLoS Biology 18 (1), e3000586.

Amorim, C. E. G., Z. Gao, Z. Baker, J. F. Diesel, Y. B. Simons, I. S. Haque, J. Pickrell, and M. Przeworski (2017). The population genetics of human disease: The case of recessive, lethal mutations. PLoS Genetics 13 (9), e1006915.

Bataillon, T. and S. F. Bailey (2014). Effects of new mutations on fitness: insights from models and data. Annals of the New York Academy of Sciences 1320 (1), 76–92.

Blumenstiel, J. P., X. Chen, M. He, and C. M. Bergman (2014). An age-of-allele test of neutrality for transposable element insertions. Genetics 196 (2), 523–538.

Bollback, J. P., T. L. York, and R. Nielsen (2008). Estimation of 2 n es from temporal allele frequency data. Genetics 179 (1), 497–502.

Booker, T. R., B. C. Jackson, and P. D. Keightley (2017). Detecting positive selection in the genome. BMC Biology 15 (1), 1–10.

Boucher, J. I., P. Cote, J. Flynn, L. Jiang, A. Laban, P. Mishra, B. P. Roscoe, and D. N. Bolon (2014). Viewing protein fitness landscapes through a next-gen lens. Genetics 198 (2), 461–471.

Boyko, A. R., S. H. Williamson, A. R. Indap, J. D. Degenhardt, R. D. Hernandez, K. E. Lohmueller, M. D. Adams, S. Schmidt, J. J. Sninsky, S. R. Sunyaev, et al. (2008). Assessing the evolutionary impact of amino acid mutations in the human genome. PLoS Genetics 4 (5), e1000083.

Brandvain, Y. and S. I. Wright (2016). The limits of natural selection in a nonequilibrium world. Trends in Genetics 32 (4), 201–210.

Bustamante, C. D., J. Wakeley, S. Sawyer, and D. L. Hartl (2001a). Directional selection and the site-frequency spectrum. Genetics 159 (4), 1779–1788.

Bustamante, C. D., J. Wakeley, S. Sawyer, and D. L. Hartl (2001b, December). Directional Selection and the Site-Frequency Spectrum. Genetics 159 (4), 1779–1788. Publisher: Genetics Section: INVESTIGATIONS.

Coop, G. (2016, March). Does linked selection explain the narrow range of genetic diversity across species? Pages: 042598 Section: Contradictory Results.

Coop, G. and R. C. Griffiths (2004, November). Ancestral inference on gene trees under selection. Theoretical Population Biology 66 (3), 219–232.

Durvasula, A. and K. E. Lohmueller (2021). Negative selection on complex traits limits phenotype prediction accuracy between populations. The American Journal of Human Genetics 108 (4), 620–631.

Evans, S. N., Y. Shvets, and M. Slatkin (2007, February). Non-equilibrium theory of the allele frequency spectrum. Theoretical Population Biology 71 (1), 109–119.

Eyre-Walker, A. and P. D. Keightley (2007). The distribution of fitness effects of new mutations. Nature Reviews Genetics 8 (8), 610–618.

Fisher, R. A. (1930). The genetical theory of natural selection: a complete variorum edition. Oxford University Press.

Fowler, D. M., C. L. Araya, S. J. Fleishman, E. H. Kellogg, J. J. Stephany, D. Baker, and S. Fields (2010). High-resolution mapping of protein sequence-function relationships. Nature Methods 7 (9), 741–746.

Gazal, S., H. K. Finucane, N. A. Furlotte, P.-R. Loh, P. F. Palamara, X. Liu, A. Schoech, B. Bulik-Sullivan, B. M. Neale, A. Gusev, and A. L. Price (2017, October). Linkage disequilibrium–dependent architecture of human complex traits shows action of negative selection. Nature Genetics 49 (10), 1421–1427.

Gazal, S., P.-R. Loh, H. K. Finucane, A. Ganna, A. Schoech, S. Sunyaev, and A. L. Price (2018, November). Functional architecture of low-frequency variants highlights strength of negative selection across coding and non-coding annotations. Nature Genetics 50 (11), 1600–1607. Bandiera abtest: a Cg type: Nature Research Journals Number: 11 Primary atype: Research Publisher: Nature Publishing Group Subject term: Functional genomics;Genome-wide association studies;Population genetics;Statistics Subject term id: functional-genomics;genome-wideassociation-studies;population-genetics;statistics.

Gravel, S. (2016). When is selection effective? Genetics 203 (1), 451–462.

Griffiths, R. (2003). The frequency spectrum of a mutation, and its age, in a general diffusion model. Theoretical population biology 64 (2), 241–251.

Griffiths, R. C. and S. Tavaré (1998). The age of a mutation in a general coalescent tree. Stochastic Models 14 (1-2), 273–295.

Haldane, J. B. S. (1932). The time of action of genes, and its bearing on some evolutionary problems. The American Naturalist 66 (702), 5–24.

Halligan, D. L. and P. D. Keightley (2009). Spontaneous mutation accumulation studies in evolutionary genetics. Annual Review of Ecology, Evolution, and Systematics 40, 151–172.

Hejase, H. A., N. Dukler, and A. Siepel (2020). From summary statistics to gene trees: methods for inferring positive selection. Trends in Genetics 36 (4), 243–258.

Hietpas, R. T., J. D. Jensen, and D. N. Bolon (2011). Experimental illumination of a fitness landscape. Proceedings of the National Academy of Sciences 108 (19), 7896–7901.

Horvath, R., M. Menon, M. Stitzer, and J. Ross-Ibarra (2022). Controlling for variable transposition rate with an age-adjusted site frequency spectrum. Genome Biology and Evolution 14 (2), evac016.

Hudson, R. R. (2002). Generating samples under a wright–fisher neutral model of genetic variation. Bioinformatics 18 (2), 337–338.

Hujoel, M. L., S. Gazal, F. Hormozdiari, B. Van De Geijn, and A. L. Price (2019). Disease heritability enrichment of regulatory elements is concentrated in elements with ancient sequence age and conserved function across species. The American Journal of Human Genetics 104 (4), 611–624.

Irving-Pease, E. K., A. Refoyo-Martínez, A. Ingason, A. Pearson, A. Fischer, W. Barrie, K.-G. Sjögren, A. S. Halgren, R. Macleod, F. Demeter, R. A. Henriksen, T. Vimala, H. McColl, A. Vaughn, A. J. Stern, L. Speidel, G. Scorrano, A. Ramsøe, A. J. Schork, A. Rosengren, L. Zhao, K. Kristiansen, P. H. Sudmant, D. J. Lawson, R. Durbin, T. Korneliussen, T. Werge, M. E. Allentoft, M. Sikora, R. Nielsen, F. Racimo, and E. Willerslev (2022, October). The Selection Landscape and Genetic Legacy of Ancient Eurasians. Pages: 2022.09.22.509027 Section: New Results.

Johri, P., B. Charlesworth, and J. D. Jensen (2020, May). Toward an Evolutionarily Appropriate Null Model: Jointly Inferring Demography and Purifying Selection. Genetics 215 (1), 173–192.

Johri, P., A. Eyre-Walker, R. N. Gutenkunst, K. E. Lohmueller, and J. D. Jensen (2022). On the prospect of achieving accurate joint estimation of selection with population history. Genome Biology and Evolution 14 (7), evac088.

Jouganous, J., W. Long, A. P. Ragsdale, and S. Gravel (2017). Inferring the joint demo-graphic history of multiple populations: beyond the diffusion approximation. Genetics 206 (3), 1549–1567.

Kamm, J. A., J. Terhorst, and Y. S. Song (2017). Efficient computation of the joint sample frequency spectra for multiple populations. Journal of Computational and Graphical Statistics 26 (1), 182–194.

Kanai, M., J. C. Ulirsch, J. Karjalainen, M. Kurki, K. J. Karczewski, E. Fauman, Q. S. Wang, H. Jacobs, F. Aguet, K. G. Ardlie, et al. (2021). Insights from complex trait fine-mapping across diverse populations. MedRxiv, 2021–09.

Kelleher, J., Y. Wong, A. W. Wohns, C. Fadil, P. K. Albers, and G. McVean (2019). Inferring whole-genome histories in large population datasets. Nature genetics 51 (9), 1330–1338.

Kichaev, G., G. Bhatia, P.-R. Loh, S. Gazal, K. Burch, M. K. Freund, A. Schoech, B. Pasaniuc, and A. L. Price (2019). Leveraging polygenic functional enrichment to improve gwas power. The American Journal of Human Genetics 104 (1), 65–75.

Kiezun, A., S. L. Pulit, L. C. Francioli, F. van Dijk, M. Swertz, D. I. Boomsma, C. M. van Duijn, P. E. Slagboom, G. van Ommen, C. Wijmenga, et al. (2013). Deleterious alleles in the human genome are on average younger than neutral alleles of the same frequency. PLoS Genetics 9 (2), e1003301.

Kim, B. Y., C. D. Huber, and K. E. Lohmueller (2017). Inference of the distribution of selection coefficients for new nonsynonymous mutations using large samples. Genetics 206 (1), 345–361.

Kimura, M. and T. Ohta (1973). The age of a neutral mutant persisting in a finite population. Genetics 75 (1), 199–212.

Lehmann, E. L. and G. Casella (2006). Theory of point estimation. Springer Science & Business Media.

Lewanski, A. L., M. C. Grundler, and G. S. Bradburd (2024). The era of the ARG: An introduction to ancestral recombination graphs and their significance in empirical evolutionary genomics. PloS Genetics 20 (1), e1011110.

Malaspinas, A.-S., O. Malaspinas, S. N. Evans, and M. Slatkin (2012). Estimating allele age and selection coefficient from time-serial data. Genetics 192 (2), 599–607.

Maruyama, T. (1974). The age of an allele in a finite population. Genetics Research 23 (2), 137–143.

Mathieson, I. and G. McVean (2013). Estimating selection coefficients in spatially structured populations from time series data of allele frequencies. Genetics 193 (3), 973–984.

Mathieson, I. and J. Terhorst (2022). Direct detection of natural selection in bronze age britain. Genome Research 32 (11-12), 2057–2067.

McVicker, G., D. Gordon, C. Davis, and P. Green (2009, May). Widespread Genomic Signatures of Natural Selection in Hominid Evolution. PLOS Genetics 5 (5), e1000471. Publisher: Public Library of Science.

Nait Saada, J., Z. Tsangalidou, M. Stricker, and P. F. Palamara (2023). Inference of coalescence times and variant ages using convolutional neural networks. Molecular Biology and Evolution 40 (10), msad211.

Neuhauser, C. and S. M. Krone (1997, February). The Genealogy of Samples in Models With Selection. Genetics 145 (2), 519–534.

O’Connor, L. J., A. P. Schoech, F. Hormozdiari, S. Gazal, N. Patterson, and A. L. Price (2019). Extreme polygenicity of complex traits is explained by negative selection. The American Journal of Human Genetics 105 (3), 456–476.

Ortega-Del Vecchyo, D., K. E. Lohmueller, and J. Novembre (2022). Haplotype-based inference of the distribution of fitness effects. Genetics 220 (4), iyac002.

Ortega-Del Vecchyo, D., C. D. Marsden, K. E. Lohmueller, et al. (2016). PReFerSim: fast simulation of demography and selection under the Poisson Random Field model. Bioinformatics 32 (22), 3516–3518.

Rannala, B. (1997). On the genealogy of a rare allele. Theoretical population biology 52 (3), 216–223.

Rasmussen, M. D., M. J. Hubisz, I. Gronau, and A. Siepel (2014, May). Genome-Wide Inference of Ancestral Recombination Graphs. PLOS Genetics 10 (5), e1004342. Publisher: Public Library of Science.

Sawyer, S. A. and D. L. Hartl (1992). Population genetics of polymorphism and divergence. Genetics 132 (4), 1161–1176.

Schraiber, J. G., R. C. Griffiths, and S. N. Evans (2013). Analysis and rejection sampling of Wright–Fisher diffusion bridges. Theoretical population biology 89, 64–74.

Shi, H., S. Gazal, M. Kanai, E. M. Koch, A. P. Schoech, K. M. Siewert, S. S. Kim, Y. Luo, T. Amariuta, H. Huang, et al. (2021). Population-specific causal disease effect sizes in functionally important regions impacted by selection. Nature communications 12 (1), 1098.

Simons, Y. B., H. Mostafavi, C. J. Smith, J. K. Pritchard, and G. Sella (2022). Simple scaling laws control the genetic architectures of human complex traits. bioRxiv, 2022–10.

Simons, Y. B. and G. Sella (2016, December). The impact of recent population history on the deleterious mutation load in humans and close evolutionary relatives. Current Opinion in Genetics & Development 41, 150–158.

Slade, P. F. (2000). Simulation of selected genealogies. Theoretical Population Biology 57 (1), 35–49.

Slatkin, M. (2000). Allele age and a test for selection on rare alleles. Philosophical Transactions of the Royal Society of London. Series B: Biological Sciences 355 (1403), 1663–1668.

Slatkin, M. (2001). Simulating genealogies of selected alleles in a population of variable size. Genetics Research 78 (1), 49–57.

Slatkin, M. and B. Rannala (2000). Estimating allele age. Annual review of genomics and human genetics 1 (1), 225–249.

Song, Y. S. and M. Steinrücken (2012). A simple method for finding explicit analytic transition densities of diffusion processes with general diploid selection. Genetics 190 (3), 1117–1129.

Speidel, L., M. Forest, S. Shi, and S. R. Myers (2019). A method for genome-wide genealogy estimation for thousands of samples. Nature Genetics 51 (9), 1321–1329.

Spence, J. P., T. Zeng, H. Mostafavi, and J. K. Pritchard (2023). Scaling the discrete-time wright–fisher model to biobank-scale datasets. Genetics 225 (3), iyad168.

Stephens, M. and P. Donnelly (2003). Ancestral inference in population genetics models with selection (with discussion). Australian & New Zealand Journal of Statistics 45 (4), 395–430.

Stern, A. J., P. R. Wilton, and R. Nielsen (2019). An approximate full-likelihood method for inferring selection and allele frequency trajectories from dna sequence data. PLoS Genetics 15 (9), e1008384.

Tennessen, J. A., A. W. Bigham, T. D. O’Connor, W. Fu, E. E. Kenny, S. Gravel, S. McGee, R. Do, X. Liu, G. Jun, et al. (2012). Evolution and functional impact of rare coding variation from deep sequencing of human exomes. Science 337 (6090), 64–69.

Vaughn, A. and R. Nielsen (2023). Fast and accurate estimation of selection coefficients and allele histories from ancient and modern dna. bioRxiv, 2023–12.

Virtanen, P., R. Gommers, T. E. Oliphant, M. Haberland, T. Reddy, D. Cournapeau, E. Burovski, P. Peterson, W. Weckesser, J. Bright, S. J. van der Walt, M. Brett, J. Wilson, K. J. Millman, N. Mayorov, A. R. J. Nelson, E. Jones, R. Kern, E. Larson, C. J. Carey, İ. Polat, Y. Feng, E. W. Moore, J. VanderPlas, D. Laxalde, J. Perktold, R. Cimrman, I. Henriksen, E. A. Quintero, C. R. Harris, A. M. Archibald, A. H. Ribeiro, F. Pedregosa, P. van Mulbregt, and SciPy 1.0 Contributors (2020). SciPy 1.0: Fundamental Algorithms for Scientific Computing in Python. Nature Methods 17, 261–272.

Williamson, S. H., R. Hernandez, A. Fledel-Alon, L. Zhu, R. Nielsen, and C. D. Bustamante (2005). Simultaneous inference of selection and population growth from patterns of variation in the human genome. Proceedings of the National Academy of Sciences 102 (22), 7882–7887.

Wiuf, C. (2001). Rare alleles and selection. Theoretical Population Biology 59 (4), 287–296.

Wohns, A. W., Y. Wong, B. Jeffery, A. Akbari, S. Mallick, R. Pinhasi, N. Patterson, D. Reich, J. Kelleher, and G. McVean (2022). A unified genealogy of modern and ancient genomes. Science 375 (6583), eabi8264.

Wright, S. (1931). Evolution in mendelian populations. Genetics 16 (2), 97.

Wright, S. (1938a, July). The Distribution of Gene Frequencies Under Irreversible Mutation. Proceedings of the National Academy of Sciences of the United States of America 24 (7), 253–259.

Wright, S. (1938b). The distribution of gene frequencies under irreversible mutation. Proceedings of the National Academy of Sciences of the United States of America 24 (7), 253.

Zeng, J., A. Xue, L. Jiang, L. R. Lloyd-Jones, Y. Wu, H. Wang, Z. Zheng, L. Yengo, K. E. Kemper, M. E. Goddard, N. R. Wray, P. M. Visscher, and J. Yang (2021, February). Widespread signatures of natural selection across human complex traits and functional genomic categories. Nature Communications 12 (1), 1164. Bandiera abtest: a Cc license type: cc by Cg type: Nature Research Journals Number: 1 Primary atype: Research Publisher: Nature Publishing Group Subject term: Genetic variation;Genome-wide association studies;Quantitative trait Subject term id: genetic-variation;genome-wide-association-studies;quantitative-trait.

Zeng, T., J. P. Spence, H. Mostafavi, and J. K. Pritchard (2023, May). Bayesian estimation of gene constraint from an evolutionary model with gene features. Pages: 2023.05.19.541520 Section: New Results.

