## Supplementary Material for "Allele ages provide limited information about the strength of negative selection"

### Supplemental Material for "Allele ages provide limited information about the strength of negative selection"

Running title: DFE estimation using allele age

### S1 An alternative method to compute the age distribution

Here, we present an alternative numerical approach to computing the densities on allele age. This approach provides us with the same quantity as the approach in the main text but differs in its implementation and speed. The main difference between the two is that the alternative approach described below computes the density on age for all segregating sample frequencies at the same time, whereas the main approach does so for a single sample frequency at a time. So, in aggregate, the alternative approach described below is faster if we want the density on ages for all sample frequencies, but the approach outlined in the main text is faster if we only want this quantity for one or a limited number of sample frequencies.

#### The density on age is proportional to the acSFS

This alternative approach is based on first computing the age conditioned site frequency spectrum—the expected number of sites present at sample frequency $i$, conditional on having arisen $a$ generations ago—or the "acSFS", across all ages in a single pass. For a diploid sample of size $n$, we denote the acSFS in the present-day (generation 0) for alleles that arose $a$ generation ago as $\Phi_{2n}^{0|a}$. Thus, $\Phi_{2n}^{0|a}[i]$ gives the expected number of alleles $a$ generations old that are found at frequency $i$ in a diploid sample of size $n$. Allowing for the possibility that the $\Phi_{2n}^{0|a}$ might be derived from a population in which the selection coefficient, population size, or mutation rate has not been constant over time, the probability that an allele is $a$ generations old given that it's found at frequency $i$ in the present is

$$P\left(a \mid i, n, \{s_0, \ldots, s_{T_{\max}}\}, \{N_0, \ldots, N_{T_{\max}}\}, \{\mu_0, \ldots, \mu_{T_{\max}}\}\right) = \frac{\Phi_{2n}^{0|a}[i]}{\Phi_{2n}^{0}[i]}, \tag{S1}$$

where $\Phi_{2n}^{0}[i] = \sum_{t=1}^{T_{\max}} \Phi_{2n}^{0|t}[i]$ is the $i^{th}$. We note that $\Phi_{2n}^{0|t}[i]$ is equivalent to $m_t[i]$ in the main text.

#### Computing the acSFS

##### Naive method

We can also compute the acSFS efficiently by iterating the same set of discrete time recursion equations for the SFS that we use in the main text approach (Jouganous et al. 2017), given the appropriate initial condition. First, let $\Phi_{2n}^{a|a}$ be the expected SFS in diploid sample of size $n$ of the mutations arising in generation $a$, assuming that the sample is taken in generation $a$. Concretely:

$$\Phi_{2n}^{a|a}[i] = \begin{cases} 2n\mu_a, & \text{if } i = 1 \\ 0, & \text{otherwise,} \end{cases} \tag{S2}$$

representing the fact that mutations arising in generation $a$ can only be present in the sample in generation $a$ as singletons, and that we expect there to be $2n\mu_a$ of them.

Next, let $\Phi_{2n}^{t|a}$ denote the expected SFS of alleles arising in generation $a$, for a sample taken in generation $t < a$ (closer to the present). To obtain $\Phi_{2n}^{0|a}$, we iterate forward in time as in the main text, but with no mutational input, i.e.,

$$\Phi_{2n}^{t-1|a} = (\mathrm{I} + \Xi_{2n,t})\Phi_{2n}^{t|a} \tag{S3}$$

starting from $t = a$ and proceeding until $t = 0$. The distribution of ages can then be obtained for all frequencies by applying Equation (S1) for each choice of $i$. In a naive implementation, we would initialize with Equation (S2) and then iterate using Equation (S3) for each possible choice of $a$ separately up to $T_{\max}$, leading to an $\mathcal{O}(nT_{\max}^2)$ runtime. In practice, we found that this method is very slow, because $T_{\max}$ is generally very large.

#### Equilibrium case

In the special case where the selection coefficient, population size, and mutation rate are constant across time, additional time-savings are possible, as we can compute all of the $\Phi_{2n}^{0|a}$ in a single pass. Key to this is realizing that we can initialize $\Phi_{2n}^{0|0}$, the SFS of *de novo* mutations arising in the current generation, as

$$\Phi_{2n}^{0|0}(i) = \begin{cases} 2n\mu, & \text{if } i = 1 \\ 0, & \text{otherwise,} \end{cases} \tag{S4}$$

and then obtain all of the remaining $\Phi_{2n}^{0|a}$ by iterating

$$\Phi_{2n}^{0|a} = (\mathrm{I} + \Xi_{2n,t})\Phi_{2n}^{0|a-1}, \tag{S5}$$

from $a = 1$ until we reach $a = T_{\max}$. This reduces the runtime to $\mathcal{O}(nT_{\max})$.

#### Non-equilibrium case

In the non-equilibrium case, we can write the calculation of the acSFS for alleles that arose in generation $a$ as

$$\begin{aligned} \Phi_{2n}^{0|a} &= \Lambda^a \times \Phi_{2n}^{a|a} \\ &= 2n\mu_a \Lambda^a[,1] \end{aligned} \tag{S6}$$

where

$$\Lambda^a = \prod_{t=1}^{a} (\mathrm{I} + \Xi_{2n,t}), \tag{S7}$$

and $\Lambda^a[,1]$ denotes the first column of $\Lambda^a$. Thus, we can compute the full acSFS in a single pass through the time axis by initializing

$$\Lambda^0 = \mathrm{I}$$

and then iterating

$$\Lambda^a = \Lambda^{a-1} \times (\mathrm{I} + \Xi_{2n,a}), \tag{S8}$$

from $a = 1$ to $a = T_{\max}$ recording $\Phi_{2n}^{0|a}$ in each generation using equation (S6). Because $\Lambda^a$ quickly becomes dense after a few iterations, each additional iteration of equation (S8) is $\mathcal{O}(n^2)$, so the total runtime is $\mathcal{O}(n^2 T_{\max})$. Notably, as long as $n$ is modest in size, we found that this algorithm was substantially faster than the naive algorithm, despite having worse scaling in $n$.

### S2 Evolution of the SFS

Here, we provide the equations to evolve the expected SFS from one generation to the next, as originally formulated by Jouganous et al. (2017). We reproduce these equations here, with slight modifications in the notation and presentation, for completeness sake. We then link the specific operators used by Jouganous et al. (2017) to the transition matrix we use in the main text.

The expected SFS at generation $t$ is represented by $\Phi_{2n}^t[.]$ (a quantity of size $2n-1$), with $n$ being the number of diploid individuals in the sample at generation $t$ in the past. The effective population size in generation $t$ is $N_t$, the selection coefficient is $s_t$ and the mutation rate is $\mu_t$. We note that while we restrict ourselves to the additive case here, Jouganous et al. (2017) give expressions for general diploid selection, so it would be straightforward to extend our method to this case. Jouganous et al. (2017) write the transition for a single sample frequency $i$ as

$$\Phi_{2n}^{t-1}[i] = \Phi_{2n}^t[i] + \frac{1}{4N_t}\tilde{\Delta}_i \Phi_{2n}^t + s_t \tilde{\nabla}_i \Phi_{2n+1}^t + 2n\mu\delta_{i=1}, \tag{S9}$$

where $\tilde{\Delta}$ and $\tilde{\nabla}$ are both sparse matrices of size $(2n-1)\times(2n-1)$ and $(2n-1)\times 2n$, respectively representing the drift and selection operators, which control the diffusion of mass to and from adjacent sample frequency bins. The subscript $i$ indicates the $i^{th}$ row, the coefficients of which are given by

$$\tilde{\Delta}_i = [\ldots,\ 0,\ \underbrace{(i-1)(2n-i+1)}_{(i-1)^{th} \text{ if } i\geq 2},\ \underbrace{-2i(2n-i)}_{i^{th} \text{ if } 1\leq i\leq 2n-1},\ \underbrace{2n-i-1}_{(i+1)^{th} \text{ if } i\leq 2n-2},\ 0,\ \ldots], \tag{S10}$$

$$\tilde{\nabla}_i = [\ldots,\ 0,\ \underbrace{i(2n+1-i)}_{i^{th} \text{ if } 1\leq i\leq 2n-1},\ \underbrace{-(2n-i)(i+1)}_{(i+1)^{th} \text{ if } 1\leq i\leq 2n-2},\ 0,\ \ldots]. \tag{S11}$$

The terms in the underbraces represent the non-zero column indices in the $i^{th}$ row (with the terms being zero if the condition isn't met), so we see that the matrices $\tilde{\Delta}$ and $\tilde{\nabla}$ are tri-diagonal and bi-diagonal, respectively.

Notably, in Equation (S9) the selection term acts on an expected SFS in a sample of size one larger than the $2n$ in our actual sample. This stems from the fact that each transmission of the less fit allele is rejected due to selection with probability $s$. When this happens, we must sample a new allele at random from the population to replace it. As a result, to resolve all possible outcomes in generation $t$ for a sample of size $2n$, we need to know the expected SFS in generation $t+1$ for a sample of size $2n+1$ (consistent with standard coalescent/diffusion assumptions, there is at most one selection event per generation, so at most one additional allele must be sampled).

To overcome this moment closure issue, Jouganous et al. (2017) use a jackknife extrapolation method first introduced by Gravel and National Heart, Lung, and Blood Institute (NHLBI) GO Exome Sequencing Project (2014) to represent $\Phi^t_{2n+1}$ as a linear function of $\Phi^t_{2n}$, which we also employ, and reproduce here for completeness.

To this end, the elements of $\Phi^t_{2n+1}$ can be written as linear combinations of $\Phi^t_{2n}$ using the equation below,

$$\Phi^t_{2n+1}[i] = \alpha_i \Phi^t_{2n}[i'-1] + \beta_i \Phi^t_{2n}[i'] + \gamma_i \Phi^t_{2n}[i'+1] \;\forall\; i \in [1, 2n-1] \tag{S12}$$

where $\alpha_i, \beta_i, \gamma_i$ are the jackknife coefficients, and $i'$ is chosen such that the frequency ${}^{i'}\!/_{2n}$ is as close as possible to ${}^{i}\!/_{(2n+1)}$ while satisfying the boundary conditions:

$$\begin{aligned} i' - 1 &\geq 1 \\ i' + 1 &\leq 2n - 1. \end{aligned}$$

The jackknife coefficients are:

$$\begin{aligned} \alpha_i &= \frac{Q_\alpha}{2(2n+2)(2n+3)(2n+4)}, \\ \beta_i &= \frac{Q_\beta}{(2n+2)(2n+3)(2n+4)}, \\ \gamma_i &= \frac{Q_\gamma}{2(2n+2)(2n+3)(2n+4)}, \end{aligned}$$

with

$$Q_\alpha = (2n+1)\left(4 + i^2\left(6 + 10n + 4n^2\right) - i\left(14 + 18n + 4n^2\right) - (2n+4)\left(2i\left(2n+1\right) - 2n - 5\right)i' + \left(4n^2 + 14n + 12\right)i'^2\right)$$
$$Q_\beta = (2n+1)\left(\left(i+1\right)\left(2n+2\right)\left(i\left(2n+3\right) - 2n - 6\right) - 2\left(2n+4\right)\left(i\left(2n+2\right) - 1\right)' i + \left(4n^2 + 14n + 12\right)i'^2\right)$$
$$Q_\gamma = (2n+1)\left(\left(i+1\right)\left(2n+1\right)\left(i\left(2n+3\right) - 2\right) - \left(2n+4\right)\left(2i\left(2n+2\right) + 2n + 1\right)i' + \left(4n^2 + 14n + 12\right)i'^2\right).$$

See Appendix D of Jouganous et al. (2017) for the full derivation.

Now, Equation (S9) can be rewritten as

$$\Phi^{t-1}_{2n}[i] = \Phi^t_{2n}[i] + \frac{1}{4N_t}\tilde{\Delta}_i \Phi^t_{2n} + s_t \tilde{\nabla}_i \mathcal{J}_{2n\to 2n+1}\Phi^t_{2n} + 2n\mu\delta_{i=1}, \tag{S13}$$

where $\mathcal{J}_{2n\to 2n+1}$ is a $2n \times (2n-1)$ matrix containing the jackknife coefficients outlined above.

We can now link equation (S13) to our presentation in the main text. There (as in Equation (1)) we write:

$$\Phi^{t-1}_{2n} = (\mathrm{I} + \Xi_{2n,t})\Phi^t_{2n} + 2n\mu_t\delta_{i=1},$$

where now we have,

$$\Xi_{2n,t} = \frac{1}{4N_t}\tilde{\Delta} + s_t \tilde{\nabla} \mathcal{J}_{2n \to 2n+1}.$$

The original implementation of `moments` employs a Crank-Nicholson method which uses a half-backward and half-forward Euler step to decrease instabilities with larger step sizes, whereas, we present our method using the fully forward Euler scheme.

### S3 Figures and Tables

Supplementary Equation S1: Table showing metrics of estimation accuracy for the strongest ($|\gamma| = 100$ or $|s| = 5 \times 10^{-3}$) and neutral selection coefficients in our simulations under the two different demographic histories and the two different thresholding schemes. In case of piece-wise exponential growth, we've listed the bias and variance in terms of the selection coefficient $s$, instead of $\gamma$ since the population size is not constant through time.

(a) Using all sites

| **Demography** | **Constant size** (using $\gamma$) | | | | | | **Exponential growth** (using $s$) | | | |
|---|---|---|---|---|---|---|---|---|---|---|
| | **Freq only** | | **Freq & age (true ages)** | | **Freq & age (est. ages)** | | **Freq only** | | **Freq & age (true ages)** | |
| | **Bias** | **Var** | **Bias** | **Var** | **Bias** | **Var** | **Bias** | **Var** | **Bias** | **Var** |
| $\gamma = -100$ ($s = -5 \times 10^{-3}$) | −2.1 | 51.3 | −1.2 | 32.3 | 0.51 | 45.4 | $2.3 \times 10^{-5}$ | $1.1 \times 10^{-7}$ | $2.6 \times 10^{-7}$ | $3.7 \times 10^{-8}$ |
| $\gamma = 0$ ($s = 0$) | 0.07 | 0.02 | −0.03 | 0.02 | −0.09 | 0.04 | $-7.2 \times 10^{-7}$ | $5.1 \times 10^{-11}$ | $-1.1 \times 10^{-6}$ | $5.2 \times 10^{-11}$ |
| $\gamma = 100$ ($s = 5 \times 10^{-3}$) | 32.6 | 42,800 | 0.13 | 5.8 | – | – | $-8.6 \times 10^{-4}$ | $3.2 \times 10^{-5}$ | $-1.5 \times 10^{-5}$ | $2.4 \times 10^{-8}$ |

(b) Using only sites with MAF > 2.5%

| **Demography** | **Constant size** (using $\gamma$) | | | | | |
|---|---|---|---|---|---|---|
| | **Freq only** | | **Freq & age (true ages)** | | **Freq & age (est. ages)** | |
| | **Bias** | **Var** | **Bias** | **Var** | **Bias** | **Var** |
| $\gamma = -100$ | −41.6 | 14,900 | −0.56 | 1,600 | 36.4 | 520 |
| $\gamma = 0$ | −4.44 | 55.2 | −2.41 | 48.2 | −6.49 | 220 |

### References

Adrion, J. R., C. B. Cole, N. Dukler, J. G. Galloway, A. L. Gladstein, G. Gower, C. C. Kyriazis, A. P. Ragsdale, G. Tsambos, F. Baumdicker, J. Carlson, R. A. Cartwright, A. Durvasula, I. Gronau, B. Y. Kim, P. McKenzie, P. W. Messer, E. Noskova, D. Ortega-Del Vecchyo, F. Racimo, T. J. Struck, S. Gravel, R. N. Gutenkunst, K. E. Lohmueller, P. L. Ralph, D. R. Schrider, A. Siepel, J. Kelleher, and A. D. Kern (2020, jun). A community-maintained standard library of population genetic models. *eLife 9*, e54967.

Gravel, S. and National Heart, Lung, and Blood Institute (NHLBI) GO Exome Sequencing Project (2014). Predicting discovery rates of genomic features. *Genetics 197*(2), 601–610.

Jouganous, J., W. Long, A. P. Ragsdale, and S. Gravel (2017). Inferring the joint demographic history of multiple populations: beyond the diffusion approximation. *Genetics 206*(3), 1549–1567.

Maruyama, T. (1974). The age of an allele in a finite population. *Genetics Research 23*(2), 137–143.

Ortega-Del Vecchyo, D., C. D. Marsden, K. E. Lohmueller, et al. (2016). PReFerSim: fast simulation of demography and selection under the Poisson Random Field model. *Bioinformatics 32*(22), 3516–3518.

Tennessen, J. A., A. W. Bigham, T. D. O'Connor, W. Fu, E. E. Kenny, S. Gravel, S. McGee, R. Do, X. Liu, G. Jun, et al. (2012). Evolution and functional impact of rare coding variation from deep sequencing of human exomes. *Science 337*(6090), 64–69.

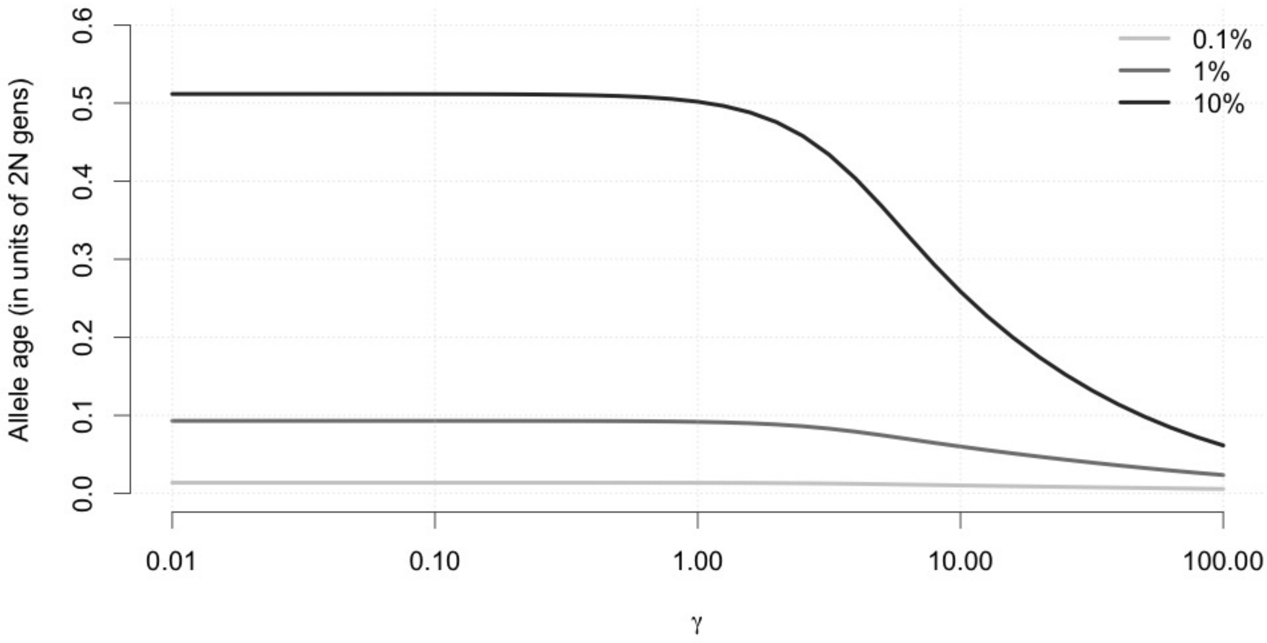


Supplementary Figure S1: Trend in average allele age under different selection strengths for a range of population allele frequencies using estimates from Maruyama (1974) for a constant population size of $N$ = 10,000. Here, we see that conditional on a particular frequency in the present day ($\{0.1, 1, 10\}\%$), the average age of the allele is much younger with stronger levels of selection. A more straightforward observation is also one in which a more frequent allele in the population will also typically be much older, for a particular selection coefficient. A dropoff is observed at $\gamma > 1$ (more apparent at higher frequencies), denoting the drift-selection boundary.

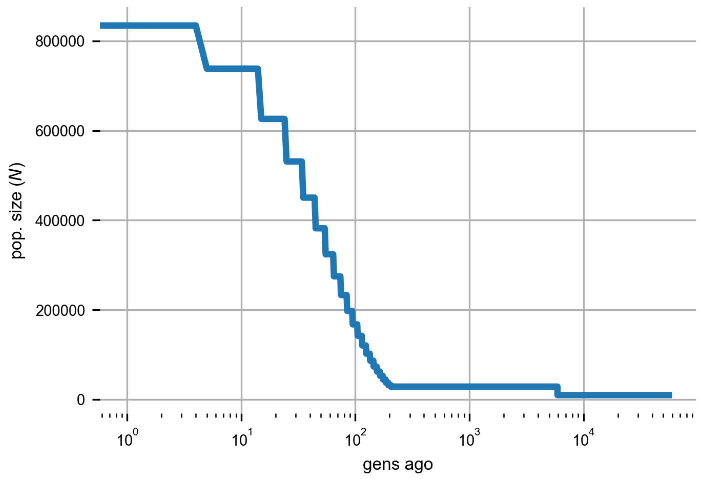


Supplementary Figure S2: The population size history for a human population inferred from $\sim$ 1,000 African-American individuals in Tennessen et al. (2012) with exponential growth in the recent past, and provided as 'Africa_1T12' in `stdpopsim` (Adrion et al. 2020). We use a discretized version of the continuous population size history for compatibility with simulation results from `PReFerSim` (Ortega-Del Vecchyo et al. 2016).

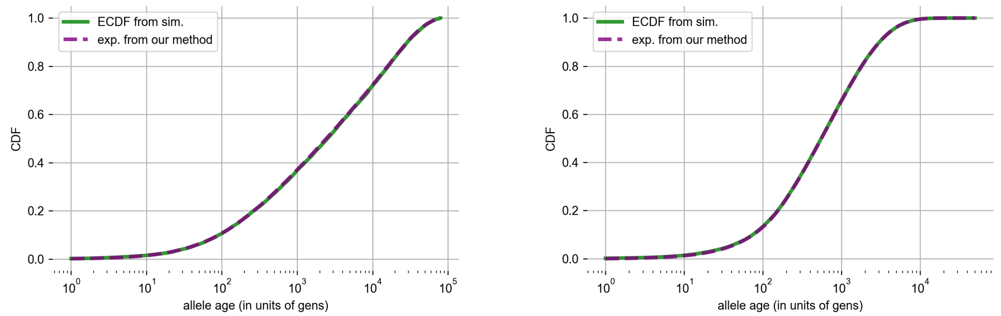


(a) For a constant population size ($N =$ 10,000) and neutrality ($s = 0$).

(b) For a human-like population history with exponential growth in the recent past and a moderate negative selection coefficient ($s = -0.0005$).

Supplementary Figure S3: Comparison between age distributions of simulations ($\sim$ 80,000 unlinked sites) from `PReFerSim` (Ortega-Del Vecchyo et al. 2016) and our approach (using a CDF) computed using the method shown in text. For both cases, **a)** constant-sized case with neutrality and **b)** human-like exponential growth case (shown in Figure S2, and inferred in Tennessen et al. 2012) with moderate selection, the cumulative distribution functions from our method overlap the distribution of simulated ages for the entire range. This indicates that our method is a fast and efficient way to computing the age distribution for any value of selection coefficient and demographic history. The Kolmogorov-Smirnov (KS) test statistic between the two distributions was found to be non-significant for the constant-sized case and 0.0065 ($p = 0.004$) for the piece-wise exponential growth case, which is a very small absolute difference.

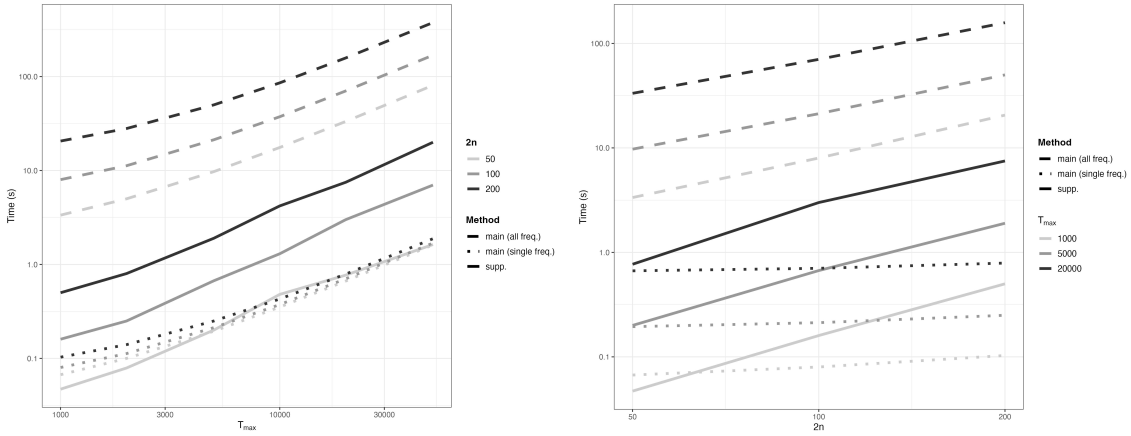


(a) Runtimes across length $T_{\max}$ (b) Runtimes across sample sizes $2n$

Supplementary Figure S4: Runtime for computing the age distribution for moderate negative selection, $s = -5 \times 10^{-4}$, and the piece-wise constant model of exponential growth from Figure S2 using the two methods listed in this manuscript: 'main (all freq.)' *in dashed* refers to the method outlined in Section 2 but summed over all sample frequencies from 1 to $2n-1$, 'main (single freq.)' *in dotted* refers to the same method but averaged over all sample frequencies, and 'supp.' *in solid* refers to the method outlined in Section S1 that necessarily computes the distribution for all sample frequencies. We observe for both methods ('main (all freq)' and 'supp.') the runtime scales linearly with both sample size, $2n$ and $T_{\max}$, which is the maximum number of generations for which the distribution on ages is calculated. The 'main (single freq.)' timing curves are simply shown here to illustrate its linear scaling properties with $T_{\max}$, but constant scaling with $2n$ (which is as expected), even though it is not a like-to-like comparison with the other two curves.

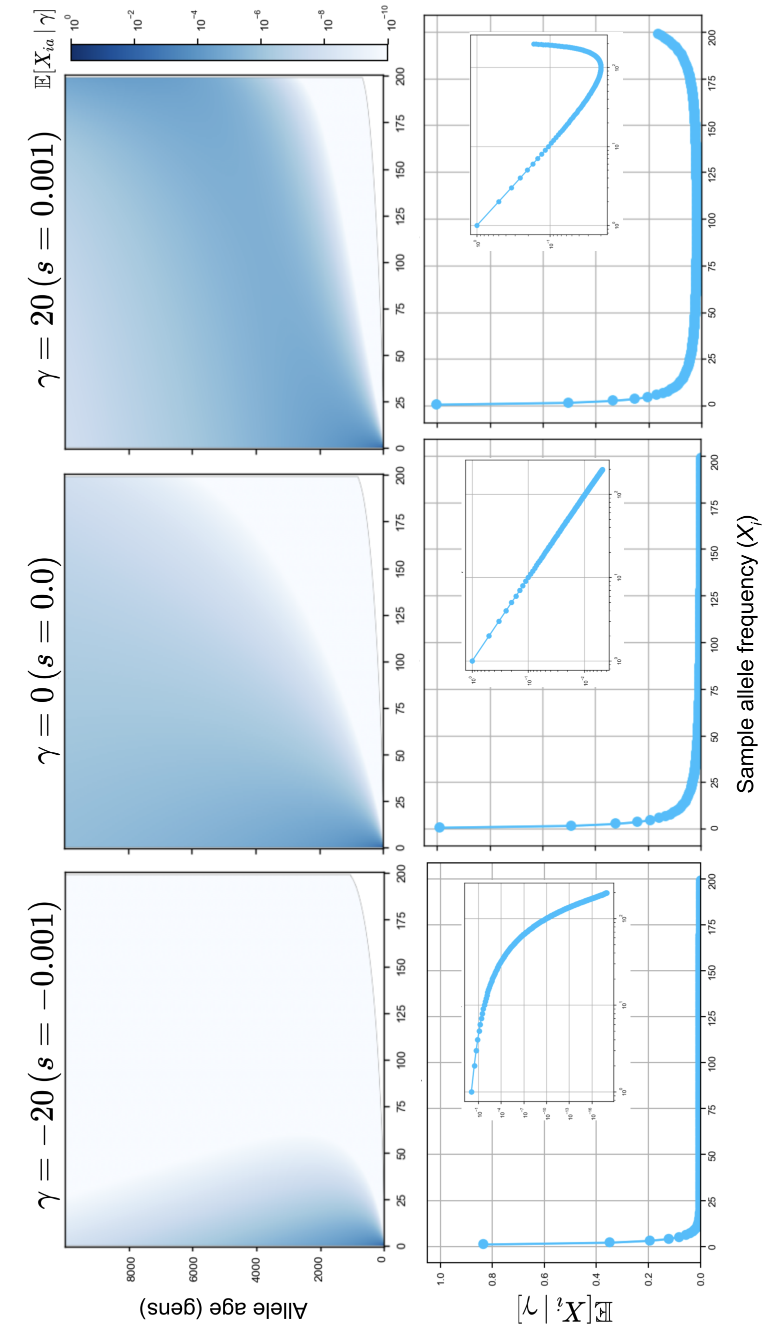


Supplementary Figure S5: Shape of expected age conditioned site frequency spectra (acSFS) and expected site frequency spectra (SFS) for different strengths of selection and constant population size ($N$ = 10,000) in a diploid sample of $n$ = 100 individuals. Overall, there is a mild correlation between frequency & age, and as the selection gets more positive, there is a larger density of alleles at higher frequencies and older ages. The shapes of the SFS in the second row will be familiar, with the insets in the second row showing the same SFS but on a log-log scale to better illustrate the deviations from neutrality.

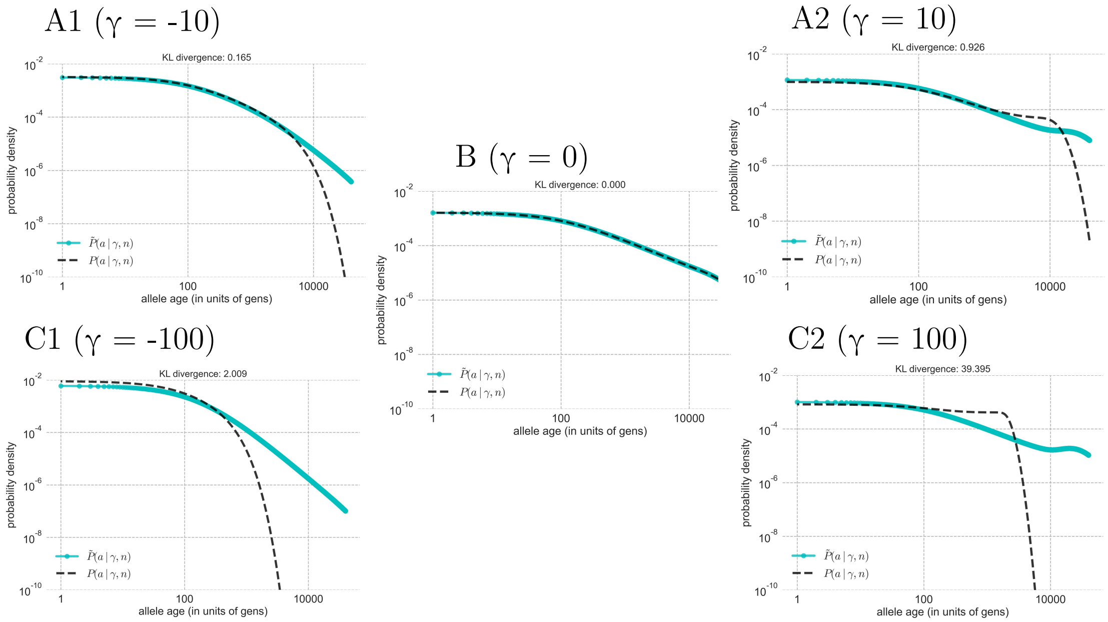


Supplementary Figure S6: The KL divergence between the unconditional age distributions of a particular selection coefficient (in black, Equation (8)) and of the approximated age distribution using the neutral frequency spectrum (in cyan, Equation (7)) across a range of selection coefficients, $\gamma$, assuming a constant population size.

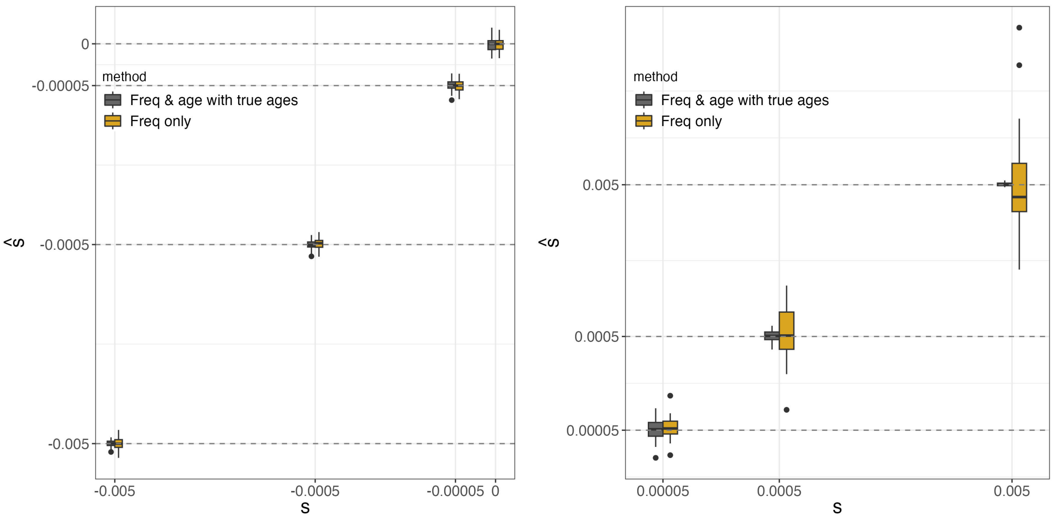


(a) For negative and neutral selection coefficients

(b) For positive selection coefficients

Supplementary Figure S7: Boxplots showing accuracy of estimation for different values of the unscaled selection coefficient, $s$, using allele frequency & age data versus allele frequency alone for a human-like demographic history. Lower and upper ends of the box correspond to first and third quartiles of the distribution of point estimates, with the whiskers extending to 1.5× the inter-quartile range (box range) with data beyond the whiskers plotted as 'outlier' points.

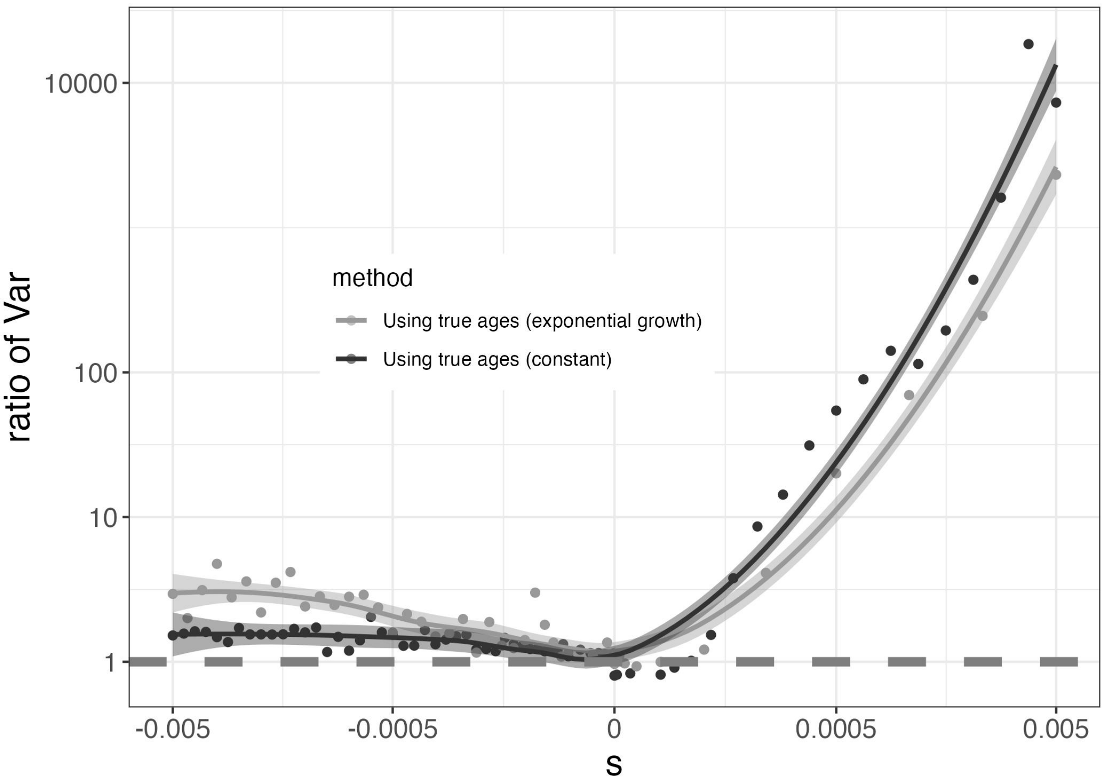


Supplementary Figure S8: Ratio of variances around the mean MLE with fitted loess lines for selection coefficients in $s \in [-0.005, 0.005]$ for both a constant population size of $N = 10{,}000$ (in black) and a piece-wise constant model of exponential growth (in gray) from Figure S2. We observe the same trend in both cases, indicating that our findings are not a consequence of the simple case of constant population size.
